# mRNA vaccines encoding membrane-anchored receptor-binding domains of SARS-CoV-2 mutants induce strong humoral responses and can overcome immune imprinting

**DOI:** 10.1101/2023.10.04.560777

**Authors:** Hareth A. Al-Wassiti, Stewart A. Fabb, Samantha L. Grimley, Ruby Kochappan, Joan K. Ho, Chinn Yi Wong, Chee Wah Tan, Thomas J. Payne, Asuka Takanashi, Horatio Sicilia, Serena L.Y. Teo, Julie McAuley, Paula Ellenberg, James P Cooney, Kathryn C. Davidson, Richard Bowen, Marc Pellegrini, Steven Rockman, Dale I. Godfrey, Terry M. Nolan, Lin-fa Wang, Georgia Deliyannis, Damian F.J. Purcell, Colin W. Pouton

## Abstract

To address the limitations of whole-spike COVID vaccines, we explored mRNA vaccines encoding membrane-anchored receptor-binding domain (RBD-TMs), each a fusion of a variant RBD, the transmembrane (TM) and cytoplasmic tail (CT) fragments of the SARS-CoV-2 spike protein. In naive mice, RBD-TM mRNA vaccines against ancestral SARS-CoV-2, Beta, Delta, Delta-plus, Kappa, Omicron BA.1 or BA.5, all induced strong humoral responses against the target RBD. Multiplex surrogate viral neutralization (sVNT) assays indicated broad neutralizing activity against a range of variant RBDs. In the setting of a heterologous boost, against the background of exposure to ancestral whole spike vaccines, sVNT studies suggested that RBD-TM vaccines were able to overcome the detrimental effects of immune imprinting. Omicron BA.1 and BA.5 RBD-TM booster vaccines induced serum antibodies with 12 and 22-fold higher neutralizing activity against the target RBD than their equivalent whole spike variants. Boosting with BA.1 or BA.5 RBD-TM provided good protection against more recent variants including XBB and XBB.1.5. Each RBD-TM mRNA is 28% of the length of its whole-spike equivalent. This advantage will enable tetravalent mRNA vaccines to be developed at well-tolerated doses of formulated mRNA.

**One Sentence Summary:** mRNA vaccines encoding membrane-anchored RBDs of SARS-CoV-2 mutants are effective vaccines that can overcome immune imprinting in mice

## INTRODUCTION

Waves of coronavirus infections resulting from the COVID-19 pandemic are caused by mutants of SARS-CoV-2 that evade humoral immunity previously acquired by way of either vaccination or viral infection. Although vaccination with an ancestral SARS-CoV-2 whole spike vaccine provides protection against serious illness (*1, 2*), boosting immunity with ancestral vaccines is ineffective at preventing infection by Omicron variants (*3–6*). To address this shortcoming, bivalent mRNA spike vaccines have been introduced encoding BA.1, or more recently BA.5, spike proteins in addition to the ancestral spike protein (*7, 8*). Unfortunately, the effectiveness of existing whole spike vaccines to prevent infection by Omicron variants appears to be compromised by the phenomenon of immune imprinting (*9, 10*), a recognised problem which can limit the effectiveness of vaccination as viruses mutate (*11, 12*). Evasive variant coronaviruses acquire mutations in the receptor-binding domain (RBD), which allow the virus to evade antibodies that bind strongly to the ancestral RBD (*13–16*). Individuals who have been exposed to the ancestral spike protein are primed to produce antibodies that bind to a range of different epitopes within this large 1273-amino acid protein. Subsequent vaccination with mRNA, encoding Omicron variants of whole spike protein, results in a significant boost in production of antibodies against ancestral epitopes, rather than focussing the immune system on induction of new antibodies that improve the ability of polyclonal antiserum to neutralize the variant RBD (*17*).

To protect ageing and vulnerable populations from future infections by evasive mutants, next generation COVID vaccines will need to overcome the problem of immune imprinting. To address this need we developed a new platform (RBD-TM) designed to focus immune responses on new antigenic epitopes resulting from mutations in the RBD. The protein coding sequences of our RBD-TM mRNA vaccines are constructed by fusing a cDNA encoding the appropriate RBD domain by way of a short spacer to cDNA encoding the ancestral transmembrane (TM) and cytoplasmic tail (CT) (Figure S1). The complete RBD-TM mRNAs have similar design features to those used in the approved whole spike vaccines (*1, 18*), and are formulated in lipid nanoparticles (LNPs) in an analogous manner.

In this manuscript we describe: (i) our initial preclinical studies comparing ancestral RBD-TM vaccine with ancestral whole spike vaccine; (ii) the development of a Beta variant (K417N, E484K, N501Y) RBD-TM vaccine which was later manufactured for human use and has undergone evaluation in a Phase 1 clinical trial; (iii) preclinical experiments in naïve mice to test immune responses to a range of variant RBD-TM vaccines, including Delta, Delta-plus, Kappa, Omicron BA.1 and BA.5 vaccines; and (iv) experiments in mice to simulate real-world vaccination against a background of exposure to ancestral SARS-CoV-2 whole spike protein. The study shows that RBD-TM mRNAs provide an adaptable platform for production of prophylactic vaccines with the potential to protect elderly and vulnerable individuals from infection by emerging coronavirus variants. A Phase 1 clinical study of the Beta RBD-TM mRNA as a fourth-dose booster vaccine has been completed. Interim data from the trial is available in preprint (*19*) and has been submitted for publication in a separate manuscript.

## RESULTS

### Immunogenicity induced by ancestral RBD-TM and whole spike mRNA vaccines

The components of the RBD-TM mRNA COVID-19 vaccine platform are shown schematically in Figures 1A-1D and are described in the methods section and in Figures S1 and S2. The RBD-TM mRNA is 28% of the length of whole spike mRNA (Figure 1A) and is designed to be expressed as a membrane-anchored monomeric RBD (Figure 1D). Unless otherwise described, this vaccine was delivered in an LNP with low content (0.15-0.25 mole%) of PEGylated lipid (Figure 1C). An initial prime and boost experiment was carried out at doses of 1, 3 or 10μg mRNA to determine whether the ancestral SARS-CoV-2 RBD-TM vaccine induced immunity in naïve BALB/c mice. Virus neutralization (VNT) studies using an early isolate of the ancestral SARS-CoV-2 virus (VIC01) or a Beta variant virus indicated that strong antibody responses were induced at doses of mRNA above 1μg (Figure 1E). Neutralization titres determined using the Beta variant virus were consistently lower at all three doses of mRNA. At the dose of 3μg, which we subsequently established is adequate for vaccination in mice, ID_50_ values were 1621 ± 644 against VIC01 and 154 ± 52 against the Beta isolate (mean ± sem, n=5). When the mice were challenged (see methods) on day 65 with VIC2089, a SARS-CoV-2 variant that had acquired the N501Y mutation in common with the Alpha variant, all three doses of ancestral RBD-TM vaccine protected the lungs from infection (Figure 1F). To compare the immune responses of mice vaccinated with either RBD-TM or whole spike mRNAs, we carried out prime and boost vaccination at either 1 or 5μg mRNA. Ancestral RBD-specific antibody titres determined three weeks after the prime or boost (i.e. day 21 or 42) are shown in Figures 1G and 1H. VNT by serum samples collected on day 42 was evaluated using VIC01 or the Beta variant (Figure 1I). At each dose the RBD-TM was more potent than whole spike although the antibody titres had reached a limiting value for either vaccine at the 5μg dose (Figure 1H). The high potency of the RBD-TM is possibly because the RBD-TM mRNA encodes 3.6-fold more RBD units than the same mass of whole spike mRNA. ID_50_ values for neutralization of VIC01 after two 5μg doses of whole spike or RBD-TM mRNA were 1277 ± 507 and 2693 ± 714 (mean ± sem, n = 5) respectively. This difference was not significant, probably a result of both vaccines being close to saturating the immune response against the target RBD at the 5μg dose. The higher activity of antisera against VIC01 after two doses of RBD-TM versus whole spike at the 1μg dose was significant (p < 0.01, t-test) as were the differences in mean VNT_50_ values against the Beta virus at either dose (p < 0.05 at both 5μg and 1μg doses, t-test). These differences were not significant when evaluated by ANOVA. The VNT studies suggested showed that a dose of 1μg whole spike mRNA failed to induce a robust neutralizing antibody response (Figure 1I). Surrogate sVNT studies were carried out in multiplex to evaluate the ability of mouse serum to inhibit of binding of a range of RBD variants to human ACE2 (Figures 1J and 1K). Details of the mutation in each RBD are shown in Figure S3. 5μg doses of whole spike vaccine induced sVNT50 values greater than 100 for all variants other than Mu and Omicron BA.1 and BA.2. The same mass of RBD-TM vaccine induced consistently higher sVNT50 values for all variants with titres above 100 for the two Omicron variants. Mean % neutralization data for four dilutions of serum samples are shown in Figure S4.

**Fig. 1.**
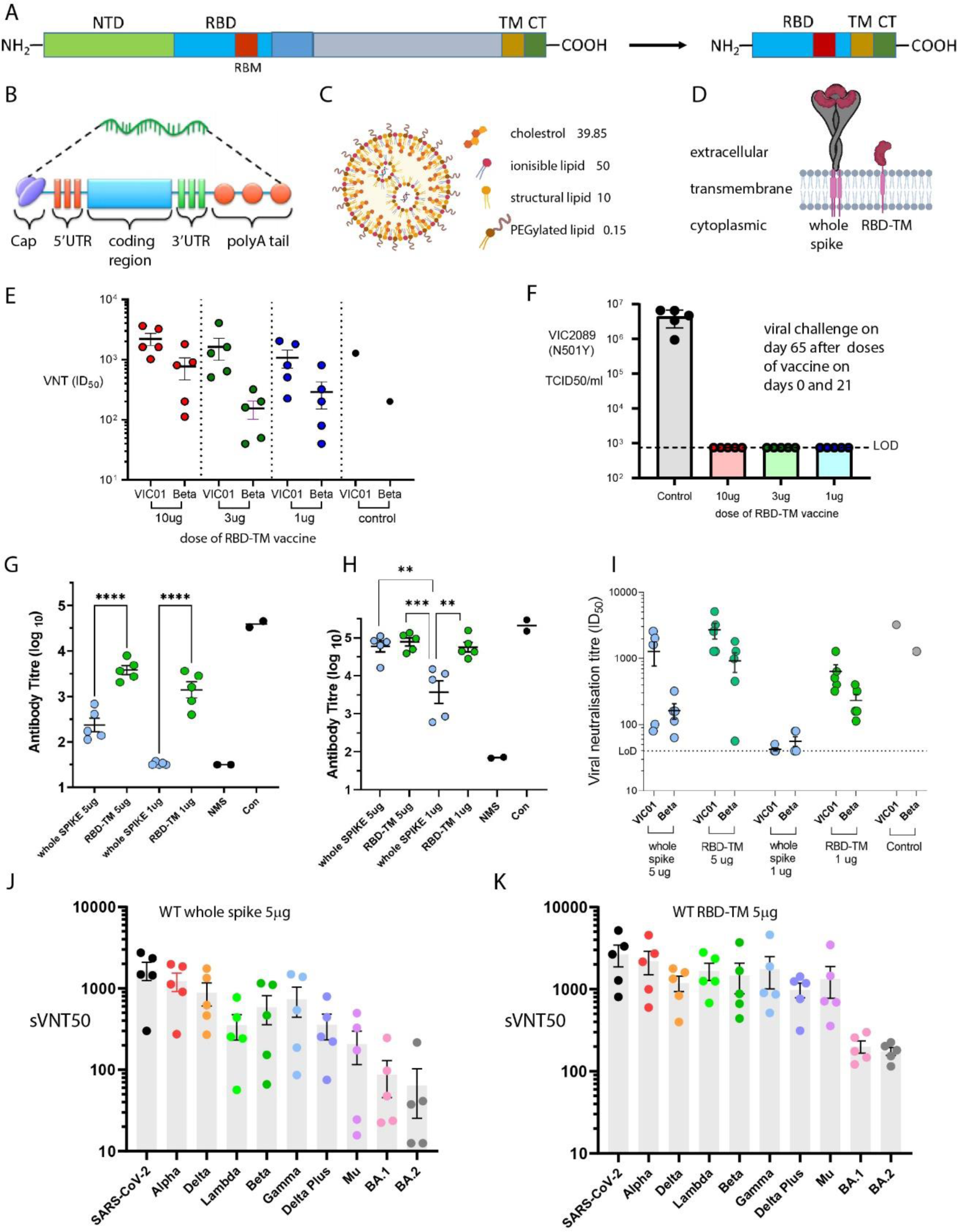
Comparison of immunogenicity induced by RBD-TM mRNA and whole spike mRNA vaccines. **(A):** Schematic diagram comparing whole SARS-CoV-2 spike protein (1273 amino acids) with the RBD-TM construct (328 amino acids). **(B):** Common features of the mRNA vaccines used in this study. We used TriLink CleanCap reagent to produce the Cap1 structure, and used a 125 nucleotide polyA tail. The UTRs were designed *de novo* with reference to known sequences. **(C):** General features of the LNP delivery system used. The identities and concentrations of ionizable and PEGylated lipids are described in the methods section. **(D):** Cartoon representation of the proteins resulting from translation of the whole-spike and RBD-TM mRNAs. **(E):** Neutralization of infection of Vero cells by serum samples from mice vaccinated intramuscularly (IM) with wild type SARS-CoV-2 RBD-TM vaccine. BALB/c mice were vaccinated on day 0 and 21 with either 1, 3 or 10μg mRNA. Serum samples were collected on day 42. Viral strains used were the VIC01 isolate of WT SARS-CoV-2 or the Beta B.1.351 variant. The control serum was obtained by pooling serum from previous vaccinations with whole spike vaccine. **(F):** Viral titres in lungs of the mice from panel (E) three days after challenge with an N501Y mutant of SARS-CoV-2 (hCoV-19/Australia/VIC2089/2020) on day 65, 44 days after the second dose of vaccine. The titre of infectious virus (TCID50; 50% tissue culture infectious dose) in the lungs of individual mice were determined by titrating lung homogenate supernatants on Vero cell monolayers and measuring viral cytopathic effect 5 days later. Control animals were unvaccinated aged-matched BALB/c mice. **(G) and (H):** RBD-specific antibody titres determined by ELISA in mouse serum samples after either 1 or 5μg doses IM on days 0 and 21 of either WT whole spike or WT RBD-TM vaccines. Antibody titres were determined on day 21(G) or on day 42 (H) **(I):** Neutralization of infection of Vero cells, by VIC01 or Beta strains of virus, by the day 42 serum samples used for ELISA studies in panel (H) (mean ± sem, n = 5). **(J) and (K):** Surrogate virus neutralization (sVNT) studies using multiplexed variant RBD-beads indicating relative neutralization of SARS-CoV-2 variants by the serum samples as used in panel (I) after doses of either 5μg of WT whole spike (J) or 5μg of WT RBD-TM (K) vaccines. Half-maximal inhibitory dilution (sVNT50) is indicated for each serum sample. Horizontal lines show mean; error bars show SEM (n = 5 mice); statistical analysis shown in G and H used ANOVA and Tukey post-hoc test, *p < 0.05, **p < 0.01, ***p < 0.001.

### Development of a Beta variant RBD-TM mRNA vaccine for clinical evaluation

In mid-2021 we began preparing for clinical evaluation of the RBD-TM platform by developing a vaccine against the Beta variant of SARS-CoV-2. Prior to the emergence of Omicron in late 2021, Beta was the VOC which had acquired the most consistent ability to evade immunity induced by ancestral vaccines (*20*). Figure 2 outlines the preclinical data supporting the development of the clinical Beta RBD-TM vaccine. Prime and boost studies were carried out in BALB/c mice vaccinated with doses between 0.1 and 10μg mRNA. Antibody titres on day 21 after the priming dose suggested that the dose-response relationship was close to linear over the range 0.1-3μg against the target Beta RBD or ancestral RBD (Figures 2A and 2B). Titres were consistently higher against the target Beta variant and could be assessed using ELISA against either Beta RBD or ancestral RBD-coated plates. Antibody titre was below 10^3^ after 0.1μg doses, even after the boost dose at day 42 or 56 (Figures 2C and 2D). Serum collected on day 56 was evaluated by VNT using ancestral VIC01 or the Beta variant (Figure 2E). A 3μg dose induced adequate protection, producing mean VNTID_50_ values of 723 ± 166 and 283 ± 53 (mean ± sem, n = 5) against Beta and VIC01 respectively. The higher activity against the target variant was not significantly different at n = 5 (p = 0.057). In a second study using a different batch of vaccine we extended the dose range to 10μg. Antibody titres and virus neutralization data at day 56 are shown in Figures 2F and 2G.

**Figure 2.**
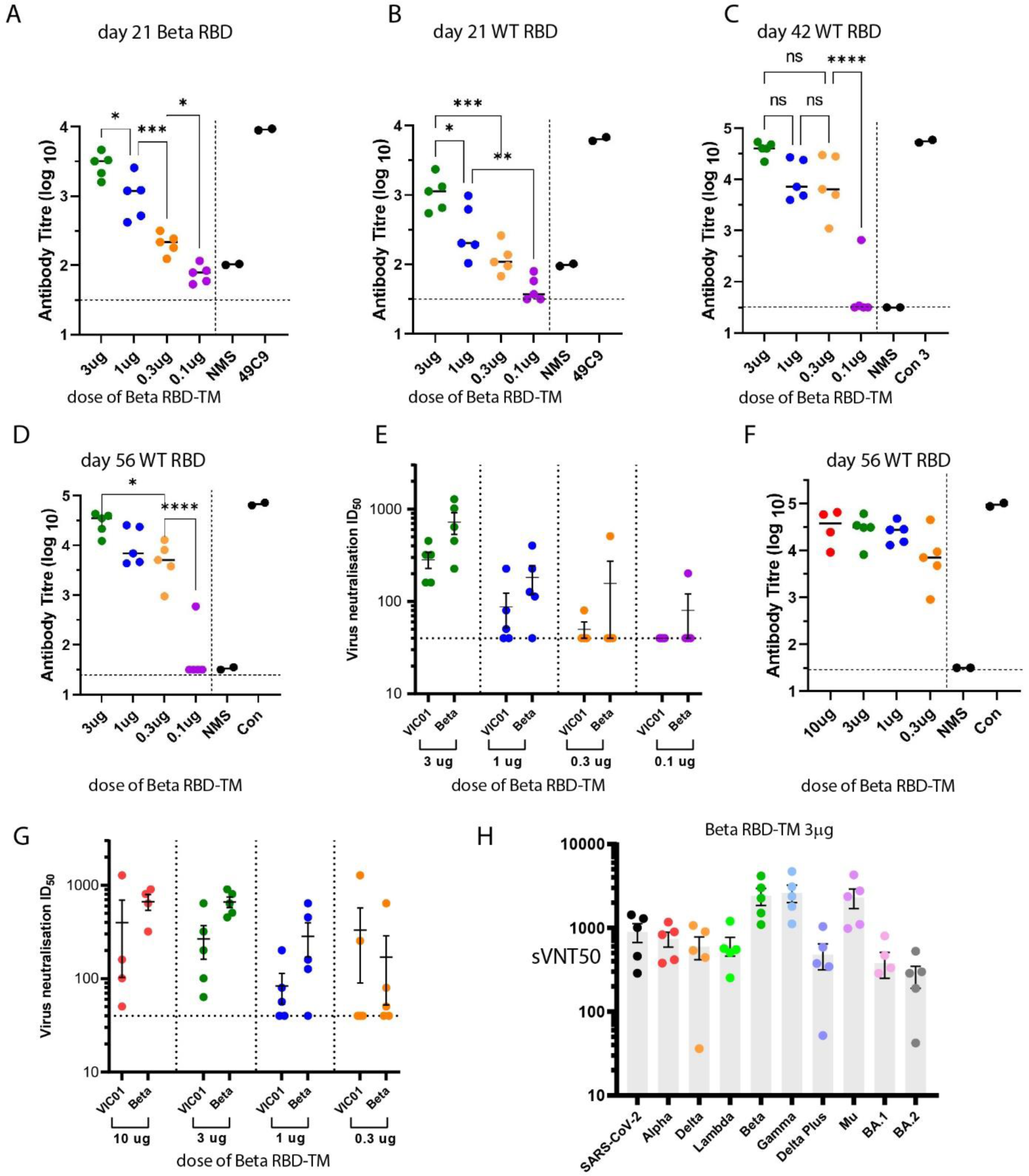
Immunogenicity of Beta RBD-TM mRNA vaccine as a function of dose. **(A)-(D):** RBD-specific antibody titres in mouse serum after doses of Beta RBD-TM vaccine administered IM on days 0 and 21. Titres on day 21 against Beta RBD (A) and titres against WT RBD on day 21 (B), day 42 (C) or day 56 (D). NMS = normal mouse serum, 49C9 = control antiserum, control = serum from mice treated with 30ug native mRNA vaccine; Horizontal lines show mean; error bars show SEM (n = 5 mice); statistical analysis ANOVA and Tukey post-hoc test, *p < 0.05, **p < 0.01, ***p < 0.001. **(E):** Neutralization of infection of Vero cells by serum samples from mice vaccinated IM with various doses of Beta RBD-TM vaccine. Neutralization of WT VIC01 or a Beta variant are shown. **(F and G):** RBD-specific antibody titres (F) and virus neutralization (G) in mouse serum at day 56 after two doses of Beta RBD-TM vaccine (day 0 and day 21) in an experiment to extend the dose range to 10μg. **(H):** sVNT study using multiplexed variant RBD-beads on day 56 after 3μg doses of Beta RBD-TM vaccine. Half-maximal inhibitory dilution (sVNT50) is indicated for each serum sample. Horizontal lines show mean; error bars show SEM (F, G, H) (n = 5 mice)

The data are in good agreement with the earlier experiments and shows that antibody responses were not enhanced by increasing the dose from 3 to 10μg. Multiplex sVNT studies indicated that ACE2 binding of all RBD variants tested was strongly inhibited by serum samples collected on day 56 after 2x 3μg doses of Beta RBD-TM, suggesting that the Beta vaccine induced broad spectrum activity. The highest sVNT50 values were observed against the Beta, Gamma and Mu variant RBDs, all of which share the E484K mutation. Inspection of mean % neutralization data at various dilutions (Figure S6), revealed that although protection against Beta, ancestral and Alpha RBDs was similar at doses of 3 and 10μg, there was a noticeable increase in inhibition of ACE2-binding by other variants, including Omicron BA.1 and BA.2 at the higher dose, suggesting that broader spectrum activity could be gained by using higher doses of RBD-TM vaccines.

In preparation for manufacture of the clinical Beta RBD-TM vaccine, we considered whether our LNP formulation would perform adequately in comparison with those used in the approved mRNA vaccines (see discussion for more detail). To evaluate this, we tested immunity induced by 3μg of our Beta RBD-TM mRNA in each of four alternative LNP formulations. The formulations differed in the choice of ionizable lipid (50 mole% MC3 or ALC-0315) and also in the mole% of PEGylated lipid used (1.5% or 0.15% DMG-PEG). Our mRNA vaccines were usually formulated using MC3 and 0.15 mole% DMG-PEG (see the discussion section). A more standard LNP formulation is MC3 with 1.5 mole% DMG-PEG. By swapping out MC3 for ALC-0315 we tested a formulation (50 mole% ALC-0315 and 1.5% DMG-PEG) which was closer to that used in the BioNTech/Pfizer COVID vaccine (*Comirnaty*), which contains 46.3 mole% ALC-0315 and 1.6 mole% of the PEGylated lipid ALC-0159. We also tested the 50 mole% ALC-0315 formulation with 0.15% DMG-PEG. A comparison of the four LNP formulations tested is shown in Figure 3. All four formulations produced strong immune responses. Antibody titres for the ALC-0315/1.5% DMG-PEG formulation were significantly higher than for the corresponding 0.15% DMG-PEG formulation when tested against ancestral RBD-coated plates (Figure 3A, 3C and 3D). The antibody titres were not significantly different to those induced by the MC3 formulations when tested against target Beta RBD-coated plates (Figure 3B and 3D). VNT studies confirmed that all four formulations induced serum samples with ID_50_ values that were not significantly different (Figure 3E). In all cases protection against viral infection was more effective against the target Beta strain. Mice were challenged with live Beta SARS-CoV-2 virus. Viral titres in the nasal turbinates or lungs were evaluated on day 3. All four vaccines protected mice from challenge by live virus (Figure 3F and 3G). To extend our evaluation of alternative formulations and comparison of the RBD-TM vaccine with whole spike vaccines, we vaccinated mice with either ancestral RBD-TM mRNA or whole spike mRNA at doses of 1 or 5μg, but this time using LNPs containing ALC-0315, cholesterol, DSPC and DMG-PEG2000 in the mole ratio 46.3: 42.7: 9.4: 1.6. This formulation differs from the formulation used for *Comirnaty* only in the identity of the PEGylated lipid. Antibody titres and VNT data are shown in Figure S5. The data confirms the higher potency of the RBD-TM versus whole spike vaccine. Both antibody titres and VNT data compared favorably with the analogous data obtained after vaccination with our MC3 50 mole%/DMG-PEG 0.15 mole% formulation (Figure 1), again giving confidence that we could prepare an RBD-TM vaccine for clinical evaluation using the latter formulation.

**Fig. 3.**
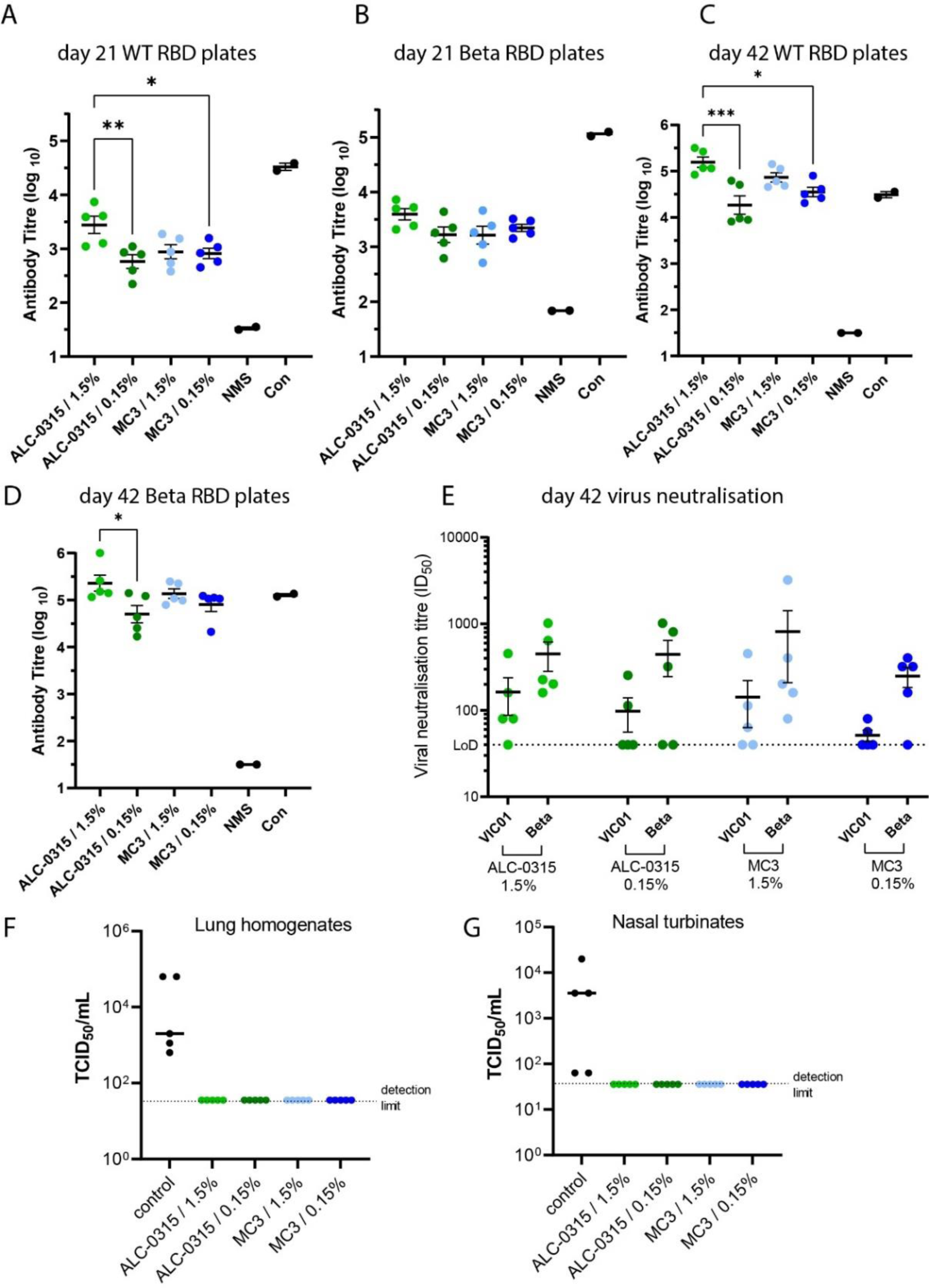
Immunogenicity and protective efficacy of the Beta RBD-TM mRNA vaccine administered in different LNP formulations. **(A-D):** WT (A, C) or Beta (B, D) RBD-specific antibody titres in Balb/c mouse serum 21 days after a single dose (A, B) or on day 42 (C,D), 21 days after a second dose of 3μg Beta RBD-TM mRNA formulated in each of four LNPs. The LNPs were formulated with either of two ionizable lipids (ALC-0315 at 46.3 mole% or DLin-MC3-DMA at 50 mole%) with either 1.5 mole% or 0.15 mole% DMG-PEG2000. Horizontal bars show mean; error bars show SEM (n = 5 mice); statistical analysis ANOVA and Tukey post-hoc test, *p < 0.05, **p < 0.01, ***p < 0.001. **(E):** Half-maximal virus neutralization titres determined against VIC01 or Beta SARS-CoV-2 using the day 42 serum samples used in C and D. Horizontal bars show mean; error bars show SEM. **(F and G):** Viral titres in lungs (F) or nasal turbinates (G) of the mice from panels (A-E) three days after aerosol challenge with a Beta variant (B.1.351) of SARS-CoV-2 on day 65, 44 days after the second dose of vaccine. Control mice were untreated aged-matched Balb/c mice. Horizontal lines show median titres in control mice. Titres in all four groups of vaccinated mice were below the limit of detection. Results reported in Figures 1, 2, 4, 5 and 6 were obtained after administration of vaccines in a single LNP formulation using 50 mole% DLin-MC3-DMA and 0.15% DMG-PEG2000.

### Activity of the Beta RBD-TM mRNA vaccine in Syrian hamsters and Sprague Dawley rats

A hamster challenge study was carried out according to the methods described in the Supplementary data file (Figure S7). This study was carried out in parallel with the evaluation of a protein Beta RBD-Fc adjuvanted by coadministration with MF59 (*21*). At doses of 3, 10 or 30μg, the RBD-TM vaccine was unable to induce full protection against infection with either ancestral or Beta SARS-CoV-2. A marginal reduction in viral titres in oropharyngeal swabs was observed but not in hamster lung tissue. With the protein Beta RBD-Fc vaccine we observed partial protection in hamsters rather than the complete protection seen in mice. These data suggest that hamsters may not respond well to RBD vaccines, on observation supported by Zhang et al (*22*) . A toxicity study to support the development of the clinical Beta RBD-TM vaccine was carried out in Sprague-Dawley rats by an independent contract research organization (Figure S7). For toxicity purposes the rats received three doses of 50μg Beta RBD-TM mRNA which was later used as the highest dose in the clinical study (*19*). High mean antibody titres greater than 10^5^ were present in the rat serum from after three doses of RBD-TM vaccine in either male (n = 15) or female (n = 15) rats (Figure S8).

### VNTs and multiplex sVNTs reveal breadth of immunity induced by alternative RBD-TM vaccines

The RBD-TM platform can be rapidly tuned to target emerging variants with mutations in the RBD domain. We vaccinated naïve mice with a range of alternative RBD-TM vaccines targeting Delta, ‘Delta-plus’, Kappa or Omicron BA.1. Data are shown in Figure 4 and Figures S8-10. The Delta variant vaccine induced serum with a spectrum of activity against other variants that were in accordance with our expectations. Mean % neutralization was greatest against the Delta RBD, strong against ancestral and Alpha, weaker against Beta, Gamma and Mu, and weaker still against Omicron BA.1 and BA.2 (Figure S9). The Delta-plus vaccine showed tighter neutralization of early variants but was weaker than Delta against BA.1 and BA.2 (Figure S10).

**Fig. 4.**
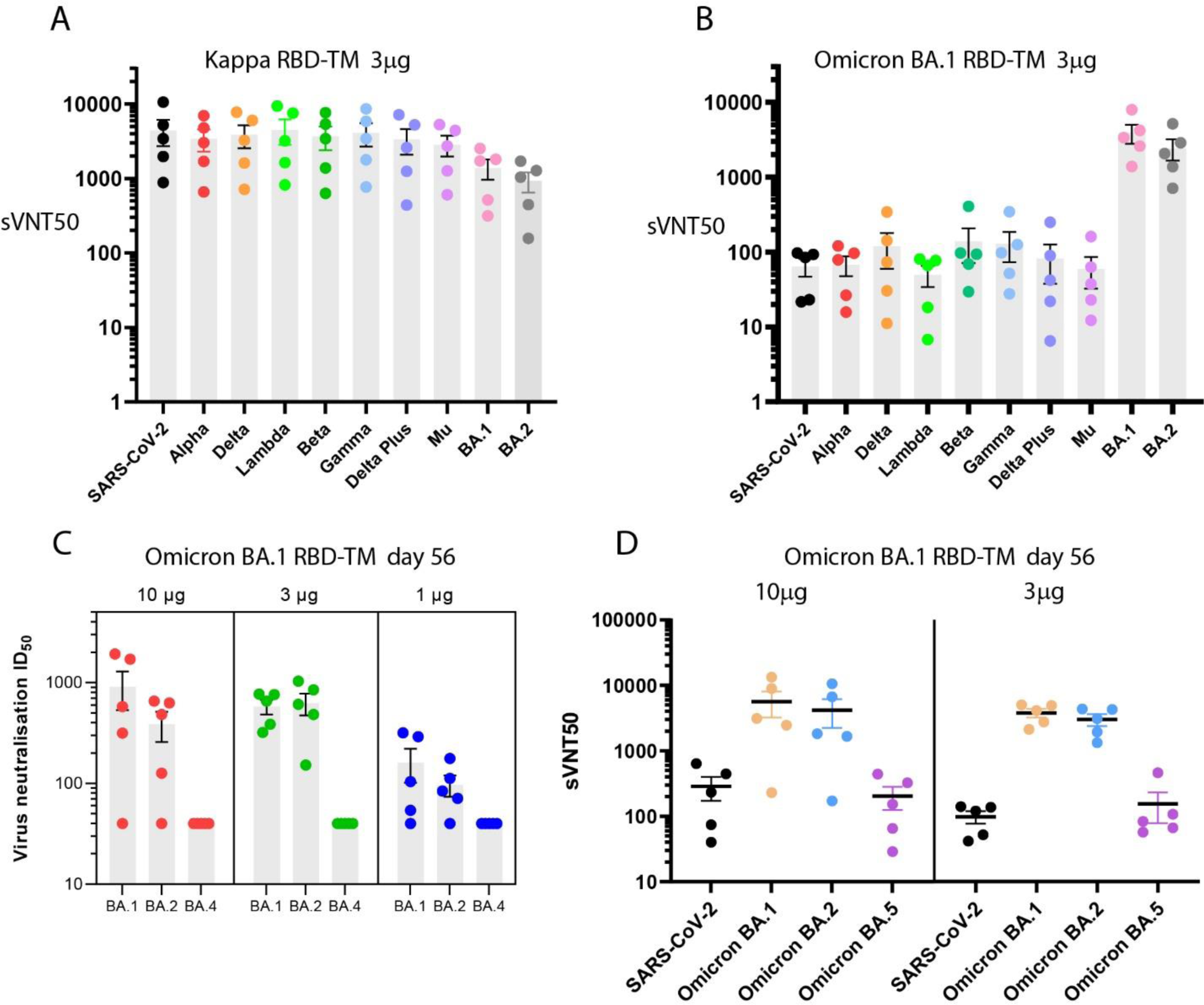
Immunogenicity of Kappa and Omicron BA.1 RBD-TM mRNA vaccines. **(A and B):** sVNT study of BALB/c mouse serum using multiplexed variant RBD-beads on day 56, after two 3μg doses (days 0 and 21) of either Kappa (A) or Omicron BA.1 (B) RBD-TM vaccines. **(C):** Half-maximal virus neutralization titres determined against naturally occurring Omicron BA.1, BA.2 and BA.4 variants of SARS-CoV-2 in serum collected on day 56 after two doses (day 0 and 21) of Omicron BA.1 RBD-TM mRNA vaccine administered at doses of 1, 3 or 10μg mRNA. BA.4 virus neutralization was below the effective limit of detection of the assay for all three doses. **(D):** sVNT study of BALB/c mouse serum on day 56 after 3 or 10μg doses of Omicron BA.1 RBD-TM mRNA using WT SARS-CoV-2, BA.1, BA.2 or BA.5-RBD-coated beads. Half-maximal inhibitory dilution (sVNT50 or VNTID50) is indicated for each serum sample. Horizontal lines or bars show mean; error bars show SEM (in all panels A-D) (n = 5 mice).

The Kappa variant RBD-TM proved to be a remarkably broad-spectrum vaccine. After two doses of Kappa vaccine, the mouse serum samples strongly inhibited ACE2 binding of all variants, including BA.1 and BA.2 (Figure 4A). In contrast the BA.1 RBD-TM vaccine produced strong inhibition of BA.1 and BA.1 RBDs but weak activity against all other earlier variants. When Omicron BA.4 and BA.5 emerged, we tested these serum samples using BA.4 virus in VNT studies (Figure 4C) and the BA.4/BA.5 RBD (which is identical) in sVNT studies (Figure 4D). The data from these assays were aligned in that the BA.1 vaccine induced serum samples that could neutralize BA.1 and BA.2 but not BA.4/BA.5. The broader neutralizing capacity of higher doses of vaccine can be observed again in the estimates of neutralizing activity of serum from mice after two doses of 3 or 10μg BA.1 RBD-TM vaccine (Figure S10). No further increases in activities after doses of 10μg were evident against BA.1 or BA.2 RBDs, but mean % neutralization values at each dilution were higher against the early pre-Omicron variants after 10μg doses versus 3μg doses. Recently, our multiplex sVNT assay was extended to include beads coupled to the RBDs of XBB, XBB.1.5 and SARS-CoV-1. The remarkable breadth of activity of the Kappa RBD-TM was further demonstrated, showing induction of strong responses even at a dose of 0.3μg against BA.5, XBB, XBB.1.5 and encouraging activity against SARS-CoV-1 at 3μg (Figure S11).

### Heterologous boost experiments in mice demonstrate that RBD-TM mRNA vaccines can overcome immune imprinting

In early heterologous boost experiments we vaccinated mice with two 5μg doses of ancestral whole spike mRNA, subsequently administering a third booster vaccine with 5μg doses of mRNA encoding either ancestral whole spike, Beta whole spike or Beta RBD-TM. The protocol for these experiments is shown in Figure 5A. Figure 5C shows that all animals had consistent antibody titres after the first and second doses of ancestral whole spike. On day 91, three weeks after the boost, there was evidence of a slight elevation in titres of Beta RBD-specific antibodies after boosting with the Beta whole spike vaccine but a significantly elevation after boost with the Beta RBD-TM vaccine. This boost was maintained at day 125 (Figure 5D). Omicron BA.1 RBD-specific antibody titre was also significantly elevated after boosting with the Beta RBD-TM. This boost may be explained in part by the 3.6-fold higher molar dose of RBD that is provided by an equal mass of RBD-TM vaccine. In more recent experiments to test boosting with Omicron variant vaccines we compared the RBD-TM vaccines as a function of mass equivalent or molar equivalent doses. As before, mice were administered doses of ancestral whole spike mRNA vaccine (8μg per dose in this experiment) on days 0 and 21. On day 56, whole spike and RBD-TM Omicron BA.1 vaccines were tested according to the booster protocol shown in Figure 5B. Group C mice received the equivalent mass of 8μg RBD-TM, group D received 2μg RBD-TM (90% of the equivalent molar dose of RBD). To investigate whether the reduced burden of LNP administered to group D could affect the immune response, we administered an additional 6μg irrelevant mRNA (encoding nanoluciferase), formulated in the same manner, to group E. The data from this experiment as assessed by sVNT is shown in Figures 5F-5K. The multiplex sVNT assay provides a rich set of data allowing comparison of neutralization titres across a range of variant RBDs. Figures 5F-5H show that enhanced immunity induced by Omicron BA.1 RBD-TM mRNA is not dependent on higher molar dose of RBD. The 2μg dose of RBD-TM was at least as effective as the 8μg dose, both of which produced elevated sVNT50 titres across the spectrum of variant RBDs.

**Figure 5.**
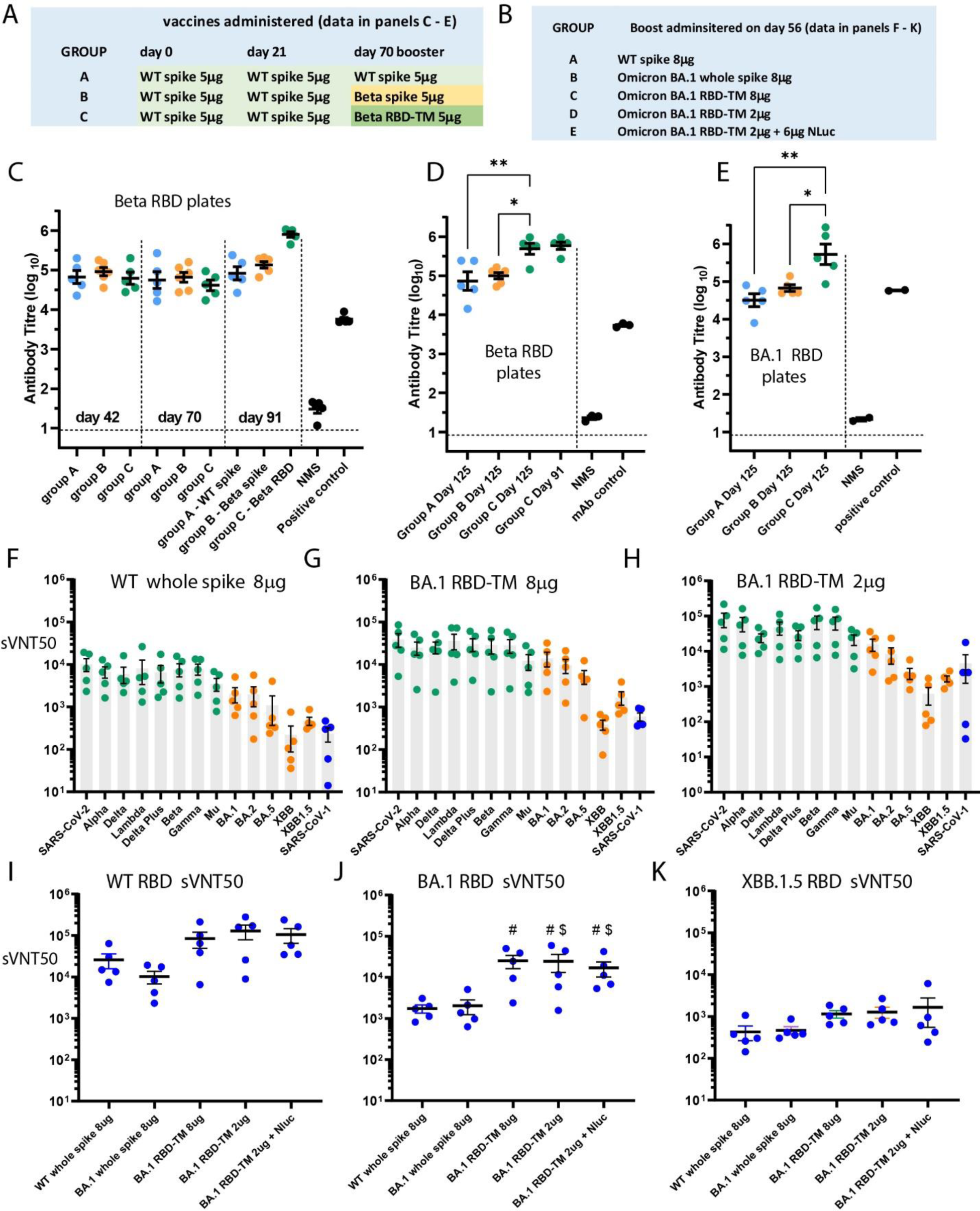
Heterologous boost studies using Beta or Omicron BA.1 RBD-TM vaccines after exposure to WT whole spike mRNA vaccine. **(A and B):** Vaccines administered to each group of mice in heterologous boost tests of Beta (A) or Omicron BA.1 (B) RBD-TM vaccines. Panel A relates to data shown in C, D and E. Panel B, which relates to the data shown in panels F-K, lists the alternative vaccines administered on day 56 after two 8μg doses of WT spike vaccine administered on days 0 and 21. **(C):** Beta RBD-specific antibody titres in BALB/c mouse serum samples at day 42 and day 70 after two 5μg doses (day 0 and 21) of WT whole spike mRNA vaccine, and on day 91 after one of three alternative booster doses administered on day 70. **(D and E):** Beta RBD-specific (D) and Omicron BA.1 RBD-specific antibody titres in mouse serum samples from the three groups on day 125. Horizontal bars show mean; error bars show SEM (n = 5 mice); statistical analysis ANOVA and Tukey pos-thoc test, *p < 0.05, **p < 0.01. **(F-H):** Half-maximal inhibitory dilution sVNT50 values determined in a multiplex RBD bead assay indicating the relative neutralization of binding of variant RBDs to ACE2 by mouse serum sampled on day 90, following day 56 boost with either 8μg WT whole spike (F), 8μg Omicron BA.1 RBD-TM (G) or 2μg Omicron BA.1 RBD-TM (H). sVNT50s for early variants, Omicron variants or SARS-CoV-1 are shown in green, orange or blue respectively. **(I-K):** Relative immunogenicity of five alternative boost vaccines administered on day 56, compared by determining sVNT50 of day 90 mouse serum samples against WT SARS-CoV-2 RBD (I), Omicron BA.1 RBD (J) Omicron XBB.1.5 RBD (K). Horizontal lines or bars show mean; error bars show SEM (in all panels F-K) (n = 5 mice). In panel J, # and $ denote vaccines that induce significantly different sVNT50 values compared to WT whole spike 8μg (#) or BA.1 whole spike 8μg ($) (p < 0.05).

In Figures 5I-5K the data is arranged to allow direct comparison of the booster vaccines against specific RBDs, i.e. ancestral, Omicron BA.1 and the recently widespread variant, XBB.1.5. Figure 5I shows that ancestral whole spike boost is more effective at neutralizing ancestral spike than the whole spike BA.1 vaccine. The RBD-TM vaccines are more effective, although in groups of 5 mice, the differences were not statistically different. When the booster effectiveness against the target BA.1 RBD are compared the differences become statistically significant by t-test (Figure 5J). It is clear that the BA.1 RBD-TM vaccine is equally effective at 8μg or 2μg with or without the 6μg LNP-encapsulated nanoluciferase mRNA.

A similar heterologous boost experiment was carried out to evaluate BA.5 vaccines using doses of 5μg mRNA in all cases. The protocol for the experiment is shown in Figure 6A. Mice received two doses of ancestral whole spike on days 0 and 21, followed by boosts with either ancestral whole spike, BA.5 whole spike or BA.5 RBD-TM on day 56. Mice were euthanized on day 70 for analysis of serum using the multiplex sVNT assay. Figures 6B-6D show sVNT50 values across the spectrum of variant RBDs. Boosting with ancestral whole spike again provided better neutralization of early SARS-CoV-2 variants than the BA.5 vaccine, but again the BA.5 RBD-TM vaccine provided higher sVNT50 titres across all variants, and was considerably more effective at neutralizing Omicron variants (shown in orange). Figures 6E-6F show the data plotted as sVNT50 titres provided by each vaccine against specific RBD variants. Figure 6E in accordance with Figure 5I demonstrates that boosting with a whole spike Omicron variant reduces neutralizing capacity against the ancestral RBD.

**Figure 6.**
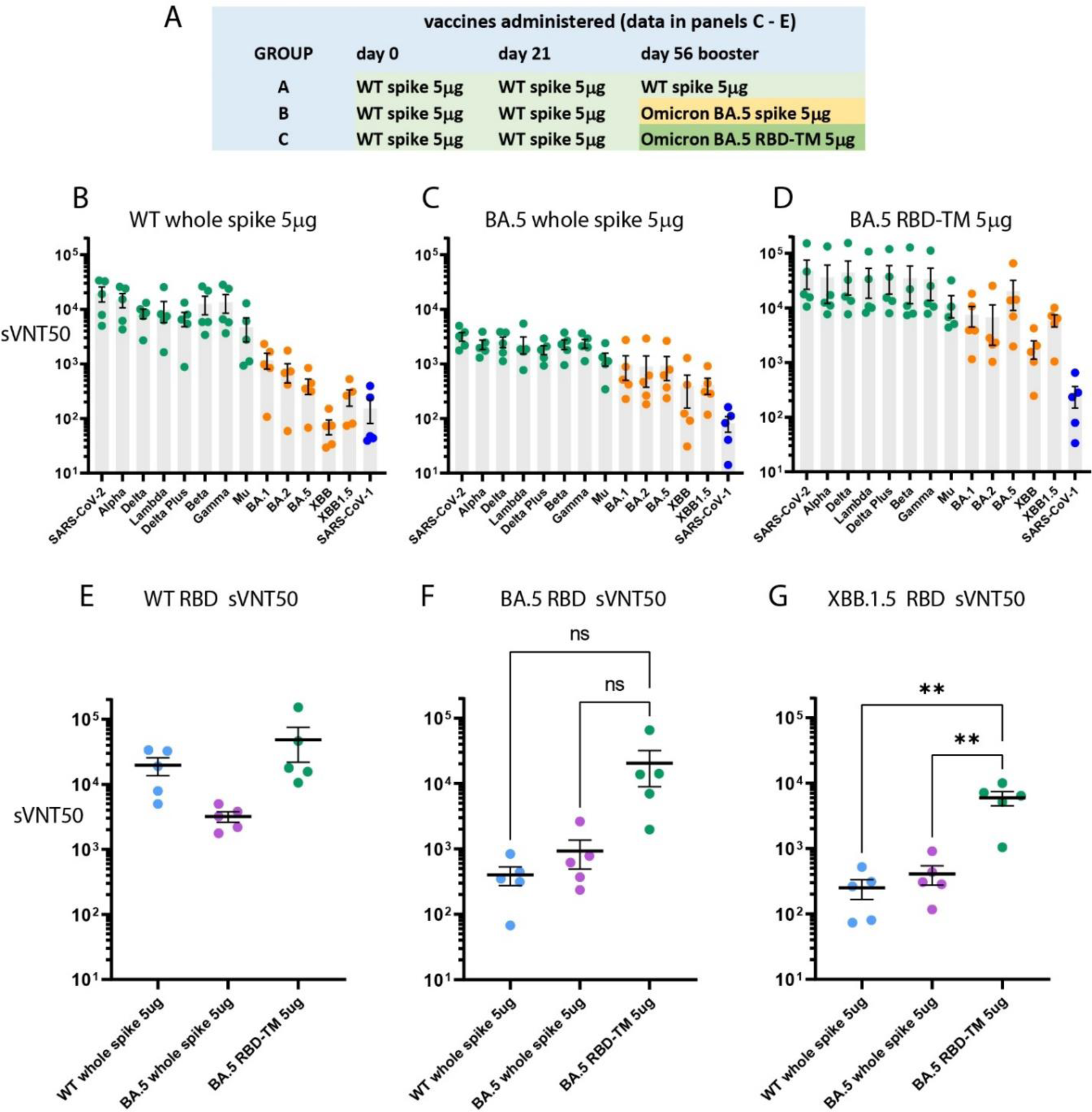
Heterologous boost study with Omicron BA.5 RBD-TM vaccine after exposure to WT whole spike mRNA vaccine. **(A):** Vaccines administered to each group of mice in the heterologous boost study shown in panels B-G. **(B-D):** Half-maximal inhibitory dilution sVNT50 values determined in a multiplex RBD bead assay indicating the relative neutralization of binding of variant RBDs to ACE2 by mouse serum sampled on day 90, following day 56 boost with either 5μg WT whole spike (B), 5μg Omicron BA.5 RBD-TM (C) or 5μg Omicron BA.5 RBD-TM (D). **(E-G):** Relative immunogenicity of three alternative boost vaccines administered on day 56, compared by determining sVNT50 of day 90 mouse serum samples against WT SARS-CoV-2 RBD (E), Omicron BA.5 RBD (F) Omicron XBB.1.5 RBD (G). Horizontal lines or bars show mean; error bars show SEM (in all panels F-K) (n = 5 mice). Statistical analysis ANOVA and Tukey post-hoc test, **p < 0.01 (no significant differences in panels E and F).

Figures 6F and 6G show that boosting with BA.5 RBD-TM is more effective at neutralizing BA.5 or XBB.1.5 RBD binding to ACE2 than either whole spike vaccine. Mean sVNT50 values obtained with BA.5 RBD-coated beads were 403 ± 126, 928 ± 436 and 20491 ± 11491 (mean ± sem, n = 5) for serum from mice vaccinated with ancestral whole spike, BA.5 whole spike or BA.5 RBD-TM respectively. The fold enhancement over ancestral whole spike was 2.3 for BA.5 whole spike and 50 for BA.5 RBD-TM. With groups of 5 animals the mean sVNT50 values for BA.5 RBD binding were not statistically different (p=0.12), but as shown by Figure 6G, the same serum samples gave statistically significant differences when compared using XBB.1.5 RBD-coated beads. Mean sVNT50 values were 250 ± 83, 411 ± 135 and 5956 ± 1465 (mean ± sem, n = 5) after vaccination with ancestral whole spike, BA.5 whole spike or BA.5 RBD-TM respectively. The fold enhancement over ancestral whole spike was 1.6 for BA.5 whole spike and 23 for BA.5 RBD-TM, emphasizing the ability of the RBD-TM platform to overcome imprinting and effectively neutralize escape variants such as XBB.1.5

## DISCUSSION

Vaccination with the ancestral whole-spike of SARS-CoV-2, and/or infection with one or more variants of the ancestral virus, have induced a level of immunity in most individuals that allows them to overcome infection by current variants of Omicron without suffering serious illness (*3-5, 7*). In contrast, during the early stages of the pandemic in 2020 and 2021, many individuals required hospitalization. The whole-spike vaccines, combined with use of face-masks and social distancing measures, were effective in preventing hospital systems becoming overwhelmed with patients that required intensive care, or at least close monitoring. In late 2023, close to three years after the onset of the COVID-19 pandemic, the mortality rate is considerably lower than it was during the first two years, but there are still many deaths occurring which are associated with COVID infection (*23*).

To reduce the incidence of COVID-related deaths, there is a need for broad-spectrum second-generation vaccines that can protect elderly and vulnerable individuals from infection by emerging variants of SARS-CoV-2 or other known betacoronaviruses. Whole-spike mRNA booster vaccines have been modified over the past year to include mutations found in the RBD of Omicron variant BA.4/5. Initially, the updated vaccines were introduced as bivalent products including the ancestral spike (*8*). The bivalent vaccines are more effective as boosters against Omicron than the ancestral spike alone (*7*), but the phenomenon of immune imprinting appears to limit the ability of variant whole-spike vaccines to induce high titres of new neutralizing antibodies against the emerging Omicron variants (*5, 6, 13, 15, 24*). The most recently updated whole-spike vaccines, which may offer more protection, target the XBB.1.5 variant, as recommended by the WHO (*25*), however other variants such as EG.5 and BA.2.86 are currently increasing in prevalence.

The RBD-TM mRNA vaccine platform offers the opportunity to develop multivalent vaccines that can be adapted to combine relevant circulating variant SARS-CoV-2 RBDs to prevent infection as the virus mutates. The RBD-TM vaccine has several advantages. Most importantly, as demonstrated in Figures 5 and 6, boosting with RBD-TM mRNAs, after two doses of ancestral whole spike vaccine, results in induction of new variant-specific antibodies, 20-50 times more effective in preventing binding of target RBDs to ACE2. Secondly, as demonstrated by Figures 1G – 1K and Figure S5, on a mass basis the RBD-TM mRNA is more potent that its whole-spike equivalent. The two vaccine platforms are equipotent on a molar basis. This implies that it will be possible to administer tri-or tetra-valent RBD-TM vaccines without exceeding the 30-50μg doses of formulated mRNA that are currently in use. Data from our Phase 1 clinical study supported this hypothesis by confirming that 10μg RBD-TM mRNA provides an effective booster dose in humans (*19*). The reactogenicity of mRNA-LNP formulations is likely to limit any increase in total dose of mRNA beyond the current dose levels. Thirdly, in the setting of a real-world heterologous boost (Figures 5 and 6), the RBD-TM induces antiserum which offers some protection against later RBD variants. For example, vaccination with 5μg BA.5 RBD-TM mRNA induced antisera with sVNT50 values greater than 1800 against the more recent Omicron variants, XBB (1825 ± 673) and XBB.1.5 (5956 ± 1465) (Figure 6D).

Experiments using the Beta RBD-TM vaccine indicated that VNTID_50_ values greater than 200-300 (Figure 3E) were adequate to achieve full protection when the same mice were challenged with live virus (Figure 3F and 3G). These data suggest that multivalent RBD-TM vaccines have the potential to protect against future escape variants. The design of multivalent vaccines is likely to improve when more data is available to help predict which RBD-TM variant vaccines have broad spectrum activity, as exemplified by the unexpected activity of the Kappa RBD-TM vaccine (Figure 4A and S10).

The precise mechanisms of action of the RBD-TM and whole spike vaccines remain to be elucidated. We hypothesized that the presentation of the whole spike protein at the surface of cells might be an important determinant of the success of the approved mRNA vaccines. Hence, we decided to include the transmembrane domain and cytoplasmic tail of the spike protein so that the translated protein would be expressed in an analogous manner as a membrane-anchored protein. We do not anticipate that RBD-TM trimers are formed, although we cannot rule out the possibility.

The tissues and cell types where protein translation occurs are critical to the induction of the immune response to whole-spike mRNA. It is known that muscle tissue translates the most protein after IM injection of mRNA formulated in LNPs, but the most important events may occur when LNPs interact with immune cells during their drainage to lymph nodes. This hypothesis is supported by our published work in which we studied the effect of LNP formulation and were able to correlate the translation of nanoluciferase reporter mRNA in lymphoid tissues with induction of immunity after IM injection of mRNA encoding ovalbumin (*26*).

We know from unpublished work that phagocytic cells (macrophages and DCs) take up LNPs and translate mRNA (*27*), but how this results in presentation to B cells or T cells is not known. We posit that, whichever cells are involved, the translation of RBD-TM mRNA occurs at the surface of the endoplasmic reticulum and the secretion signal sequence subsequently results in presentation of a plasma membrane-anchored RBD. We concluded early in our studies (Figures 1G – 1K) that, perhaps surprisingly, presentation of the RBD-TM in this way is equivalent, with respect to the molarity of RBD moieties, to presentation of RBD as a component of the trimeric whole-spike protein. This implies that the location of the RBD translated from RBD-TM mRNA and its subsequent processing by antigen-presenting cells is likely to be similar to the presentation of RBD antigens from whole-spike mRNA.

The LNP formulation used for the preclinical studies described here, and also for our Phase 1 clinical study (*19*), made use of lipids that have been previously used in FDA-approved products for human use. The ionizable lipid, DLin-MC3-DMA, is used in Onpattro (*28*), as are distearoylphosphatidylcholine (DSPC) and cholesterol. The latter two ‘structural’ or ‘helper’ lipids are also used in the Moderna mRNA vaccine, *Spikevax*, as is the neutral PEGylated lipid, DMG-PEG_2000_, which we used throughout this study (*29*).

We used an LNP formulation with some differences to the ‘standard’ Onpattro formula, which uses ionizable lipid, cholesterol, DSPC and the PEGylated lipid, PEG_2000_-C-DMG, in the following ratio; 50:38.5:10:1.5 mole% (*28*). Firstly, we typically used a formula with reduced PEGylated lipid content, ie. 50: 39.85:10:0.15 mole%. Secondly, we reduced the total lipid content, usually characterized by an N/P ratio of 6, to a N/P ratio of 5, where N/P ratio denotes the molar ratio of DLin-MC3-DMA to nucleotide, i.e. the molar ratio of ionizable nitrogen atoms to anionic phosphate moieties. These changes have the effect of producing LNPs that are typically larger (in the 120-160nm range as compared to 60-100nm range) and which are more negatively charged than standard LNPs. By considering the available surface area and availability of PEGylated lipids, we estimate that, before administration, our LNPs have a reduced mantle of PEG at the LNP-water interface, which may affect uptake by phagocytic cells. However, we recognize that the surface properties of DMG-PEG_2000_-coated LNPs will be substantially changed, by interaction with lipoproteins and plasma proteins, when they reach the blood circulation after IM injection (*30*).

The modifications in lipid molar ratios we made to the standard LNP formula were based on our previous, as yet unpublished, observations (*27, 31*). Prior to our work on the COVID vaccine we were investigating the effect of size, charge and PEGylated lipid content on biodistribution. We found that after intravenous (IV) injection, larger, more negatively charged particles are extracted to a lesser extent by the mouse liver. This results in a higher extent of mRNA translation in spleen versus liver (*27, 31*). We posit that this redistribution, and reduction in liver uptake, is desirable for vaccination, in contrast with the aim of the Onpattro product which was designed for delivery of siRNA to the liver (*28*).

The spleen/liver redistribution is evident when the PEGylated lipid content is reduced within the range 0.15-0.25 mole% PEGylated lipid and with N/P ratio = 5 (*27, 31*). We found that redistribution correlated with improved translation of reporter mRNA in phagocytic cells in the spleen and improved cell-mediated responses to vaccination of mice with mRNA encoding ovalbumin. The redistribution correlates with the observation by Siegwart and colleagues, who also showed that negative charge is associated with delivery to the spleen (*32–34*).

ELISAs and viral neutralization assays using serum from vaccinated naïve mice suggested that immunity against the target antigen was predictable, reproducible, and approximated to a linear function of dose of mRNA over the range 0.3 to 3μg. Antibody titres and VNTID_50_ titres were marginally higher at 10μg doses against the target variant, but the improvement in broad spectrum activity at higher doses was evident from the multiplex sVNT assays. This implies that design of multivalent RBD-TM vaccines will require careful analysis of the breadth of activity produced by specific mRNAs. The surprising strength and breadth of activity of the Kappa RBD-TM vaccine illustrates this point. Analysis of the data from multiplex sVNT assays (*35*) suggested that this is a precise and reliable approach for analyzing breadth of immune responses. The rank order of sVNT50 values was consistent with our expectations, based on the degree of similarity between target variant vaccine and each alternative RBD tested. When we were able to compare sVNT data with neutralization studies using viral infection of Vero cells, there was good agreement between the two assay methods. This supports data suggesting that neutralizing antibodies are predominantly bound to the RBD (*36*). Consequently, we suggest that sVNT assays using serum from vaccinated mice will provide accurate predictions of protection induced by specific RBD-TM vaccines against emerging or pre-emergent RBD variants.

Since the onset of the COVID pandemic there has been interest in the potential of RBD-based vaccines, given the proportion of neutralizing antibodies that bind to the RBD (*36*). The majority of RBD vaccine candidates in clinical development are recombinant proteins, though there have been two notable mRNA candidates. A mRNA vaccine encoding secreted RBD ^(319-541)^, ARCoV-mRNA, was developed in China by Walvax/Abogen and is now approved for use in Indonesia (*37*). Early in BioNTech/Pfizer’s clinical program, a secreted RBD fusion protein modified with the addition of a T4 fibritin-derived foldon domain, designed to form a trimeric complex (BNT162b1) was tested in a Phase 1 study (*38*). This candidate did not progress when the whole spike vaccine BNT162b2 was selected for its Phase 3 efficacy study(*2*). The RBD-TM platform described here has been evaluated in a joint Phase 1 clinical trial using the Beta variant RBD-TM in parallel with an adjuvanted Beta RBD-Fc protein vaccine (*19, 21*). Given that the ancestral vaccines offered comparatively good neutralization of the Beta variant, as compared to Omicron variants, it was not clear how well the RBD-TM vaccine was able to overcome immune imprinting in humans. A subsequent clinical efficacy study using a contemporary Omicron variant would be valuable.

How the membrane-anchored RBD-TM compares with secreted RBD vaccines in terms of mechanism of action and potency is not known but is currently under investigation. We were not aware of any other parallel developments of membrane-anchored mRNA RBD vaccines until a recent report of a Phase 1 clinical study of a self-amplifying RNA construct was published (*39*). The advantage of mRNA RBD-based vaccines is the speed with which the product can be modified as the virus mutates, and the potential for design of multivalent vaccines. The shorter RBD-TM design will allow multivalent vaccines to be produced without risking increased reactogenicity. The challenge will be to outflank the RBD mutation of the virus by anticipating the emergence of evasive mutants (*16, 40-42*), making use of the developing knowledge of likely hot-spots in the RBD, and deep scans that identify which mutations are tolerated without loss of ACE2 binding (*43–45*).

## MATERIALS AND METHODS

### Study design

The objectives of the study were to determine: (i) whether RBD-TM mRNA SARS-CoV-2 vaccines were as effective as whole spike vaccines in mice; (ii) whether RBD-TM vaccines could be tailored to vaccinate against alternative variants of SARS-CoV-2 and produce broad spectrum vaccines; and, most importantly, (iii) to determine whether RBD-TM vaccines are able to induce new antibody production against new variants against a background of immune imprinting with ancestral whole spike protein. The preclinical studies also allowed us determine a suitable range of doses for a clinical study of an RBD-TM vaccine which was carried out in parallel with ongoing preclinical experiments. We used mice for the majority of the preclinical studies but also carried out some challenge tests in Syrian hamsters and a vaccine toxicity study in rats. For comparison of immune responses to vaccines, i.e. dose response studies, comparisons between RBD-TM and whole spike vaccines, or comparisons between formulations, the experiments were usually carried out using groups of five mice for each treatment. This number of mice per group was generally adequate to establish statistical significance and served to keep animal numbers within acceptable limits as approved by Monash Institute of Pharmaceutical Sciences animal ethics committee. Data were not excluded under any circumstances. To avoid investigator bias, serum samples from mice were coded so that the laboratory scientists carrying out ELISA and neutralization tests were unaware of which serum samples were under analysis. Biological variation of immune responses within groups of animals was generally greater than between experiments comparing replicate batches of vaccine formulations. Routine replicates using different batches were not carried out.

### RBD-TM mRNA production

The mRNAs used in this study were produced using HiScribe T7 mRNA synthesis kit (NEB, Australia) using linearized DNA produced by PCR amplification. The transcribed mRNAs included a 3’-UTR with Kozak sequence and 5’-UTR, both designed de novo, and included polyA_125_ tails. The sequences were optimized to reduce the uridine content of mRNA. We used N1-methyl-pseudoUTP instead of UTP to produce chemically modified mRNA, in common with the two approved COVID-19 vaccines. CleanCap reagent AG (TriLink) was used in accordance with the manufacturer’s recommendations to produce Cap1 chemistry at the 5’ terminus. The mRNA was subject to cellulose purification before use.

### Lipid nanoparticle (LNP) formulation

The following lipids were used in the study: the ionizable lipids used for most studies was (6*Z*,9*Z*,28*Z*,31*Z*)-Heptatriaconta-6,9,28,31-tetraen-19-yl-4-(dimethylamino)butanoate (‘DLin-MC3-DMA’, MedChemExpress, USA) and for comparisons we used [(4-hydroxybutyl)-azanediyl]-di-(hexane-6,1-diyl)-bis-(2-hexyldecanoate) (ALC-0315). The commonly used helper lipids were cholesterol (Sigma-Aldrich, Germany) and 1,2-distearoyl-sn-glycero-3-phosphocholine (DSPC) (Avanti Polar Lipids Inc., USA). The PEGylated lipid used was 1,2-dimyristoyl-rac-glycero-3-methoxypolyethylene glycol-2000 (DMG-PEG 2000) (Avanti Polar Lipids Inc., USA). Formulations of the vaccines into LNPs involved the following steps: an aqueous solution of mRNA at pH4 was mixed with a solution of the four lipids in ethanol, using a microfluidics mixing device (NxGen Ignite Nanoassemblr) supplied by Precision Nanosystems. The suspension of nanoparticles was adjusted to a pH of 7.4 using a 1:3 dilution in Tris buffer, then dialysed against 25mM Tris buffer to remove the ethanol. The LNP suspension was adjusted with sucrose solution to produce the cryoprotected, isotonic final form of the product. The product was sterile filtered (0.22um) prior to being aliquoted into sterile vials for storage at −80°C. Characterization of the LNPs included analysis for RNA content, encapsulation efficiency, RNA integrity. Particle size and polydispersity index (PDI) were determined by dynamic light scattering, a standard method for submicron dispersions, using a Zetasizer (Malvern Instruments). Typically, encapsulation efficiency was 85-95%, particle size (Z-average) of the 0.15% PEGylated lipid formulation was 120-160nm with PDI < 0.2

### Intramuscular inoculation of mice

All animal experiment procedures described were conducted under the approval of the Monash Institution of Pharmaceutical Science Animal Ethics Committee. BALB/c mice (n=5 per group) were vaccinated by injection of LNP-mRNA suspension (50μl) into the calf muscle. Mice were primed on day 0 and boosted on day 21. Mice were bled just prior to the second injection, and typically three weeks (day 42) and five weeks (day 56) following the second injection. Viral challenge studies were carried out on day 65. For heterologous boost experiments BALB/c mice (n=5 per group) were vaccinated on days 0 and 21 with a mRNA-LNP vaccine produced in house encoding the whole spike of ancestral SARS-CoV-2. On day 56 the mice were vaccinated for the third time with one of a series of test booster vaccines. Mice were bled on days 21, 56 and at later times after the booster vaccine.

### Enzyme-linked immunosorbent assay (ELISA) for measurement of RBD-specific antibody responses

WT and variant RBD-specific total antibody responses in the sera of mice pre- and post-inoculation were investigated by ELISA using the RBD monomer from either the WT, Beta or Omicron BA.1 variant strain. Flat bottom 96 well maxisorp plates (ThermoFisher Scientific) were coated with 50μl/well of RBD monomer at a concentration of 2 μg/ml in Dulbecco’s phosphate buffered saline (DPBS; Gibco Life Technologies). Plates were incubated overnight at 4°C, after which unbound RBD monomer was removed, and wells were blocked with 100 μl/well of 1% bovine serum albumin (BSA fraction V, Invitrogen Corporation, Gibco) in PBS for 1-2 hours before washing with PBS containing 0.05% v/v Tween-20 (PBST). Mouse sera were added to wells and left to incubate overnight at room temperature. After washing, bound Ab was detected using horseradish peroxidase (HRP)-conjugated rabbit anti-mouse Ig Abs (Dako, Denmark). The detection antibody was incubated for 1 hour at room temperature in a humidified atmosphere and the plates then washed five times with PBS/ 0.1% Tween 20. 100μl of tetramethylbenzidine substrate (TMB, BD Biosciences) was added to each well for 5-7 minutes before the reaction was stopped using 100 μl/well of 1M orthophosphoric acid (BDH Chemicals, Australia). Optical density (OD) of each well was determined at wavelengths of 450 nm and 540 nm. Titres of Ab were expressed as the reciprocal of the highest dilution of serum required to achieve an OD of 0.3 which represents at least five times the background level of binding.

### In vitro micro-neutralisation assay (VNT)

SARS-CoV-2 isolates hCoV/Australia/VIC01/2020 (Ancestral) and hCoV/Australia/ QLD1520/2020 (Beta) were passaged in Vero cells and aliquots stored at −80 °C. Mouse serum samples were heat inactivated at 56 °C for 45 min prior to use. Serum was serially diluted in MEM medium, followed by the addition of 100 TCID_50_ of SARS-CoV-2 in MEM/0.5% BSA and incubation at room temperature for 45 min. Vero cells were washed twice with serum-free MEM before the addition of MEM containing 1 μg/ml of TPCK trypsin. Vero cells were then inoculated in quadruplicate with the plasma:virus mixture and incubated at 37 °C and 5% CO2 for 3-5 days. For SARS-CoV-2 omicron variants hCoV-19/Australia/NSW/RPAH-1933/2021 [BA.1], hCoV-19/Australia/VIC/35864/2022 [BA.2] and hCoV-19/Australia/VIC/55437/2022 [BA.4], variants were passaged in Calu3 cells in DMEM with 2% FCS and microneutralisation assays were performed in Vero E6-TMPRSS2 cells. Cytopathic effect was scored, and the neutralising antibody titer was calculated using the Reed–Muench method.

### RBD-ACE2 multiplex inhibition assay

For multiplex determination of surrogate virus neutralization titres (sVNTs), we adapted the Luminex platform as described previously (*35*). AviTag-biotinylated receptor-binding domain (RBD) proteins from different SARS-CoV-2 variants and other sarbecoviruses were coated on MagPlex-Avidin microspheres (Luminex) at 5 μg per 1 million beads. RBD-coated microspheres (600 beads per antigen) were preincubated with serum at a final dilution of 1:20 or greater for 1 hour at 37°C with 800 rpm agitation. After 1 hour of incubation, 50 μl of phycoerythrin (PE)– conjugated human angiotensin-converting enzyme 2 (ACE2) (hACE2; 1 μg per milliliter; GenScript) was added to the well and incubated for 30 minutes at 37°C with agitation, followed by two washes with 1% bovine serum albumin in phosphate-buffered saline (PBS). The final readings were acquired using a MAGPIX reader (Luminex) and expressed as half-maximal inhibitory dilution (sVNT50).

### Mouse SARS-CoV-2 challenge model

Protective efficacy against upper (nasal turbinates) and lower (lung) airways infection was assessed using a mouse SARS-CoV-2 challenge model with a human clinical isolate of SARS-CoV-2, VIC2089 (N501Y) variant (hCoV-19/Australia/VIC2089/2020) or a naturally arising Beta (K417N, E484K, N501Y) variant, B.1.351. Vaccinated and unvaccinated control mice were aerosol challenged with 1.5 x 10^7^ TCID_50_ infectious units of VIC2089 or B.1.351 using either an inhalation exposure system (Glas-col) (Fig 1) or venturi nebulisation (Fig 3). Three days later challenged mice were euthanised and infectious virus titres (TCID_50_; 50% tissue culture infectious dose) in the lungs (Figs 1 and 3) and nasal turbinates (Fig 3) of individual mice were determined by serial dilution of lung or nasal supernatants onto confluent Vero cell (clone CCL81) monolayers. Plates were incubated at 37°C for four or five days before measuring cytopathic effect under an inverted phase contrast microscope. TCID_50_ was calculated using the Spearman and Krber method.

### Statistical methods

ANOVA tests with post-hoc multiple comparisons, and Student’s t tests, were calculated using GraphPad Prism

### List of Supplementary Materials

Figure S1. Amino acid sequence of SARS-CoV-2 (wild type) ‘RBD-TM’ (328 residues)

Figure S2. Chemical structure of lipids used for LNP formulation

Figure S3. Mutations in the variant RBDs used in this study

Figure S4. sVNT analysis of mouse serum samples following vaccination of naïve mice with 5μg doses of wild type whole spike or RBD-TM vaccines

Figure S5. sVNT analysis of mouse serum samples following vaccination of naïve mice with 3μg or 10μg doses of an RBD-TM vaccine encoding the Beta variant of SARS-CoV-2

Figure S6. Comparison of the immunity induced by RBD-TM and whole spike mRNA vaccines delivered in an LNP similar to that used in the BioNTech/Pfizer product

Figure S7. Challenge study carried out in Syrian hamsters

Figure S8. Antibody titres in serum of vaccinated rats

Figure S9. sVNT analysis of mouse serum samples following vaccination of naïve mice with Delta, ‘Delta-plus’ or Kappa RBD-TM vaccines

Figure S10. sVNT analysis of mouse serum samples following vaccination of naïve mice with 3μg or 10μg doses of Omicron BA.1 RBD-TM vaccine

Figure S11. sVNT_50_ values determined using serum samples taken on day 56 from individual mice after vaccination with either 3, 1 or 0.3μg Kappa RBD-TM vaccine

## Acknowledgments

This work was supported primarily by grants from the Australian Medical Research Future Fund (MRFF), the Victorian State Government entity, mRNA Victoria. GD was supported by philanthropic funds from IFM, DIG was supported by an NHMRC Investigator Award 2008913. The following colleagues are acknowledged for their support or comments during the study: Mason Littlejohn, Jason Mackenzie, Joseph Torresi, Stephen Kent, Adam Wheatley, Sharon Lewin, David Jackson and Kanta Subbarao. Drew Brockman and Peter Tapley (Agilex Biolabs) are acknowledged for their participation in the rat toxicity studies reported in the supplementary materials. Thanks to Julian Druce and Leon Caly (Victorian Infectious Disease Reference Laboratory) for isolating and distributing ancestral and variant SARS-CoV-2 virus during the course of this study.

## Funding

Medical Research Future Fund (MRFF) Award 2005846: DIG, TMN, DFJP, GD, SR, CWP Victorian Government, mRNA Victoria: DFJP, CWP

Monash University PhD Scholarships: TP, AT, HS,

Australian National Health and Medical Research Council 2008913: DIG

Singapore National Medical Research Council (MOH-COVID19 RF-003)

IFM investors: GD

## Author contributions

Conceptualization: CWP, HAW

Methodology: HAW, SAF, CWT, JM, PE, KCD, RB, GD

Investigation: HAW, SLG, RK, JKH, SLYT, CWT, TJP, AT, HS, JM, PE, JPC, KCD, GD

Visualization: CWP, MP, SR, DIG, TMN, L-FW, DFJP

Funding acquisition: CWP, DIG, DFJP, TMN

Project administration: CWP, DIG

Supervision: CWP, RB, MP, L-FW, DFJP, GD

Writing – original draft: CWP

Writing – review & editing: All authors

## Data and materials availability

All data are available in the main text or the supplementary materials

## SUPPLEMENTARY MATERIALS

**Figure S1.**
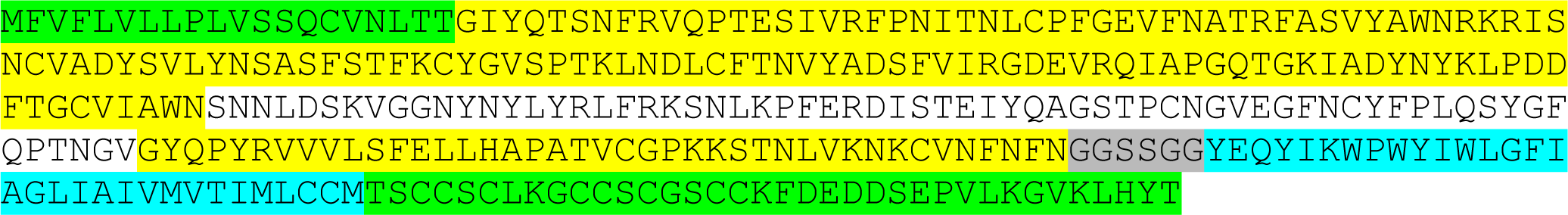
Amino acid sequence of SARS-CoV-2 (wild type) ‘RBD-TM’ (328 residues) Annotation: Signal peptide residues 1-20 (green); RBD (receptor binding domain) equivalent to residues 311-545 (yellow); RBM (receptor binding motif) equivalent to 438-506 (white); linker (GGSSGG); TM (transmembrane domain) equivalent to 1206-1237 (turquoise); CT (cytoplasmic tail) equivalent to 1238-1273 (green).

**Figure S2.**
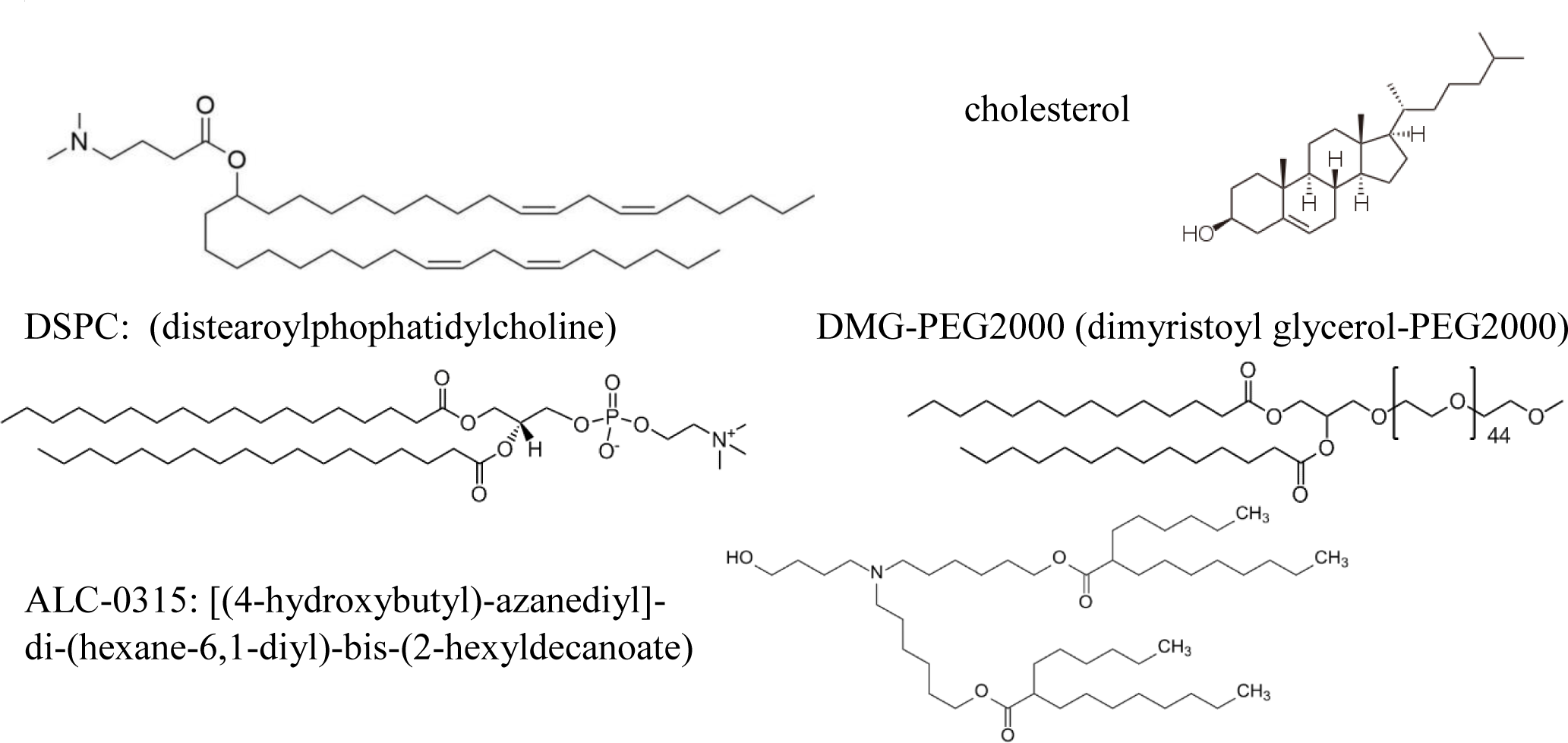
lipids used for LNP formulation. DLin-MC3-DMA: [(6Z,9Z,28Z,31Z)-heptatriaconta-6,9,28,31 tetraen-19-yl-4-(dimethylamino) butanoate]

**Figure S3:**
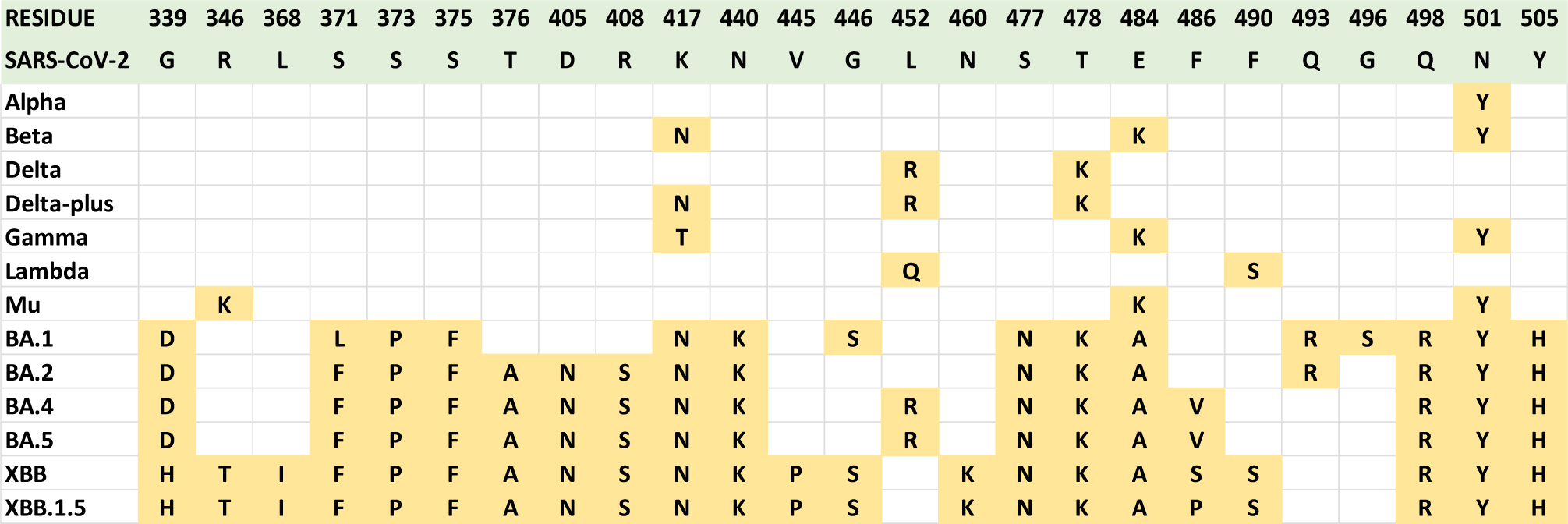
Mutations in the variant RBDs used in this study. The amino acid residue numbers and amino acid identities in the ancestral SARS-CoV-2 are shown shaded in green. The figure shows only the residues that are mutated in at least one variant RBD used in the study. Mutations are shown shaded in orange for each variant RBD. The residue numbering relates to the whole spike protein (1-1273) of SARS-CoV-2. Note that the RBD is within residues 330-528.

**Figure S4:**
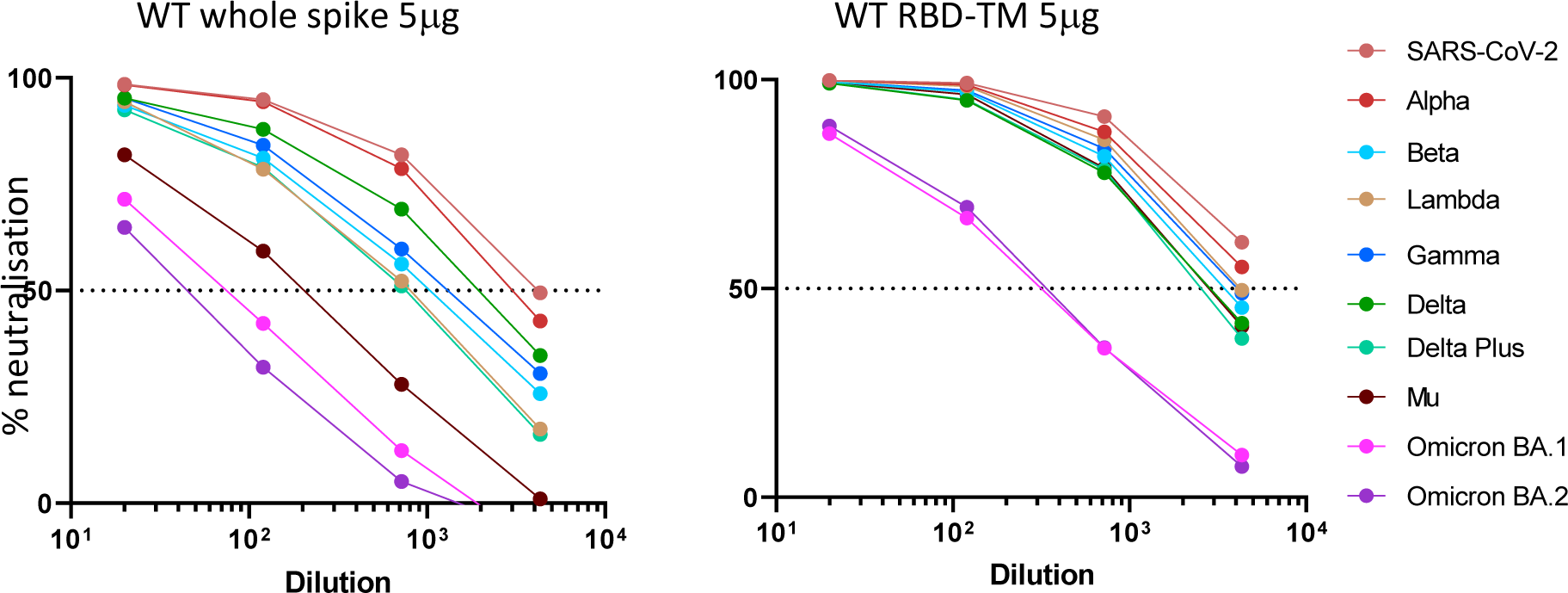
sVNT analysis of mouse serum samples following vaccination of naïve mice with 5μg doses of wild type whole spike or RBD-TM vaccines. Mice were vaccinated on day 0 and day 21. Graphs show mean % neutralisation of binding of variant RBD-beads to ACE2 plotted against dilution of the mouse serum samples collected on day 56 (n = 5 mice).

**Figure S5:**
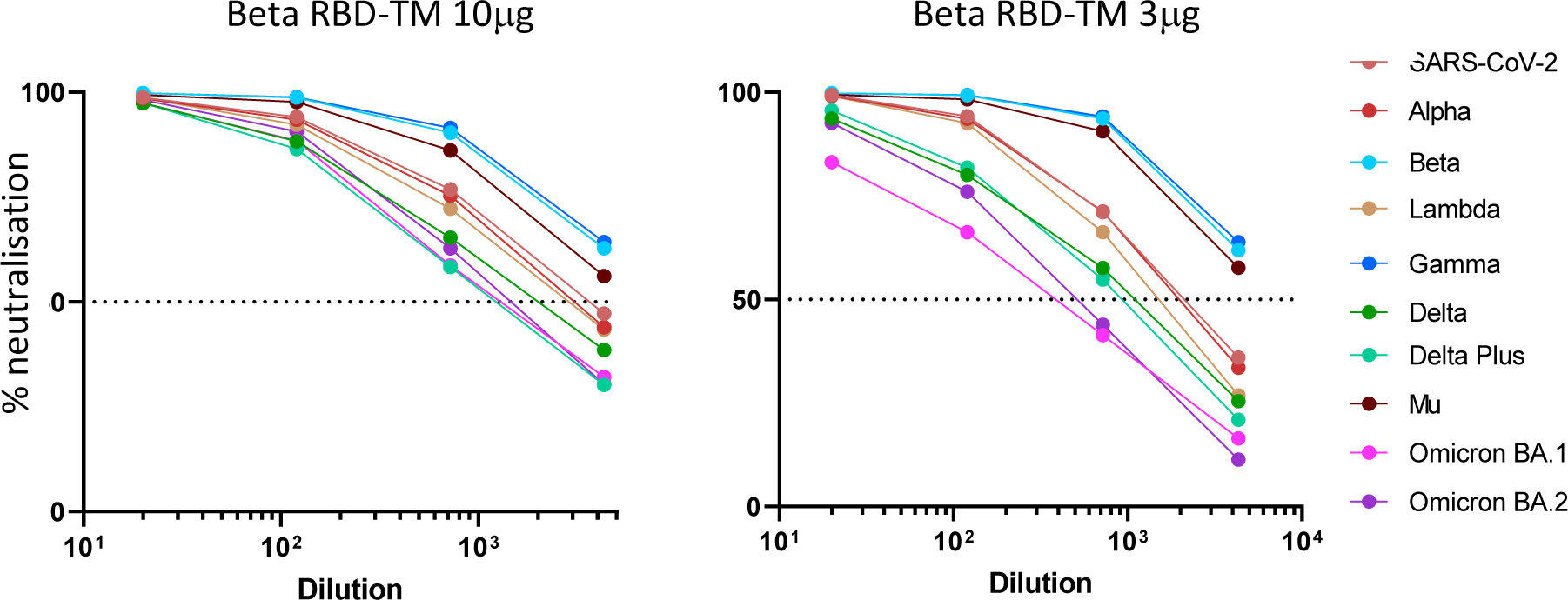
sVNT analysis of mouse serum samples following vaccination of naïve mice with 3μg or 10μg doses of an RBD-TM vaccine encoding the Beta variant of SARS-CoV-2. Mice were vaccinated on day 0 and day 21. Graphs show mean % neutralisation of binding of variant RBD-beads to ACE2 plotted against dilution of the mouse serum samples collected on day 56 (n = 4 (10μg) or 5 (3μg) mice).

**Figure S6:**
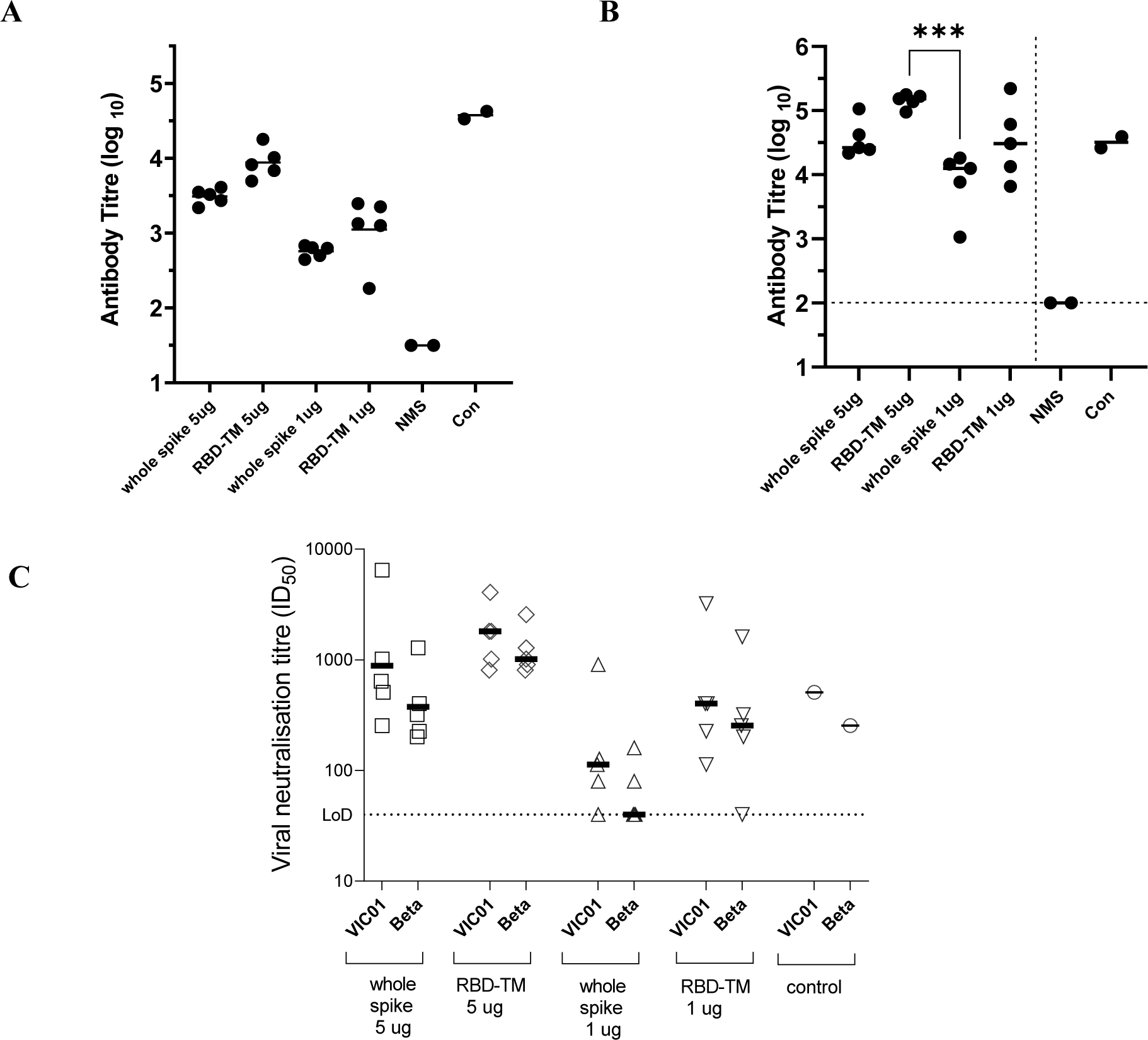
Comparison of the immunity induced by RBD-TM and whole spike mRNA vaccines delivered in an LNP similar to that used in the BioNTech/Pfizer product. Each mRNA was administered at either 1μg or 5μg on days 0 and 21. Serum samples were collected on days 21 and 42 (n = 5 mice per group). Panels A and B: Antibody titres in mouse serum samples on day 21 or day 42. (NMS = normal mouse serum, Con = in house sample of high titre mouse serum from WT vaccinated mice) Panel C: Virus neutralisation titre (ID_50_) quantifying the ability of mouse serum (day 42) to inhibit infection of Vero cells by WT (VIC01) or a Beta variant of SARS-CoV-2. (Con = Con = in house sample of high titre mouse serum from WT vaccinated mice) LNP Formulation: the LNP formulation used contained ALC-0315, cholesterol, DSPC and DMG-PEG2000 in the mole ratio 46.3: 42.7: 9.4: 1.6. This formulation differs from that used in the commercial product *Comirnaty* in that we used DMG-PEG200 rather than the PEGylated lipid ALC-0159, which was unavailable at the time when the experiment was performed.

**Figure S7.**
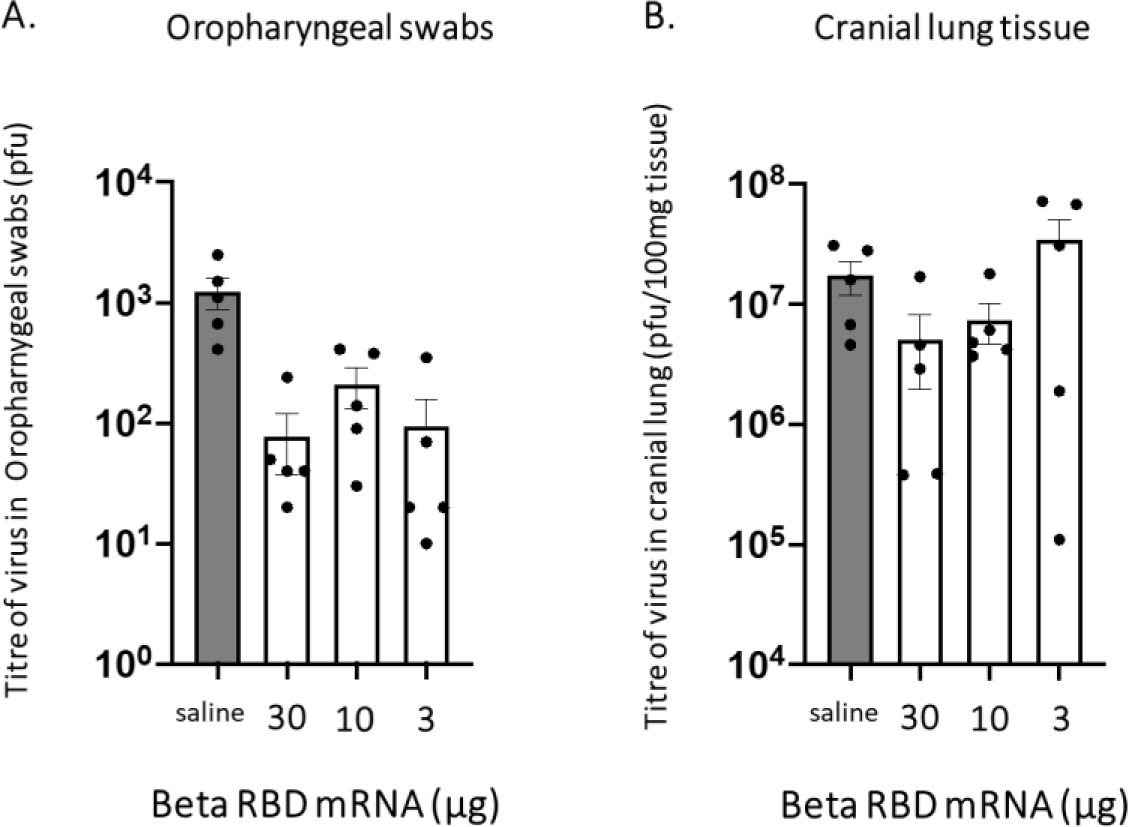
Challenge study carried out in Syrian hamsters. **Summary of the study:** At Colorado State University, protective efficacy against upper (oropharynx) and lower (lung) airways infection was assessed using a golden Syrian hamster SARS-CoV-2 challenge model with a human clinical isolate of SARS-CoV-2, WT (WA-01/USA) or a naturally arising Beta (K417N, E484K, N501Y) variant, B.1.351. These studies were conducted under approval number 1106 from the Colorado State University Institutional Animal Care and Use Committee (13Jul2020). **Methods**: Male hamsters were vaccinated with 30, 10 or 3µg of the Beta RBD-TM mRNA vaccine receiving 2 intramuscular doses on days 0 and 21. A 3rd dose was administered on day 63, and challenge performed on day 85. Hamsters were challenged under ketamine-xylazine anesthesia by intranasal instillation of 10^4^ pfu of SARS-CoV-2 virus in a volume of 100µl. Oropharygeal swabs were collected on days 1, 2, and 3 post-challenge. Three days post-challenge, hamsters were sacrificed and infectious virus titres (pfu/gram) in turbinate, cranial lung and caudal lung tissue were determined by plaque assay of homogenates on Vero cell monolayers. Neutralising antibody titres against the WT strain were assessed in sera collected 2 weeks after the third dose of vaccine using the plaque reduction neutralisation test (PRNT) using a 90% cutoff. Sample sizes of 5-10 hamsters per group were used. Humane endpoint was loss of 20% or more of challenge day body weight, which was not observed. Monitoring for activity, nasal discharge and other clinical signs was performed daily. **Results:** Reduced viral titres in oropharyngeal swabs was apparent in vaccinated mice at all three doses (Figure S6A) but the cranial lung tissue titres appeared to be unaffected (Figure S6B). Serum antibody titres were not significantly different from controls and the PRNT test suggested that serum samples were unable to protect hamsters from infection. Similar results were obtained in parallel studies of a protein vaccine (RBD-hFc dimer with MF59) which were carried out as part of the same study. These surprising data suggest that Syrian hamsters do not respond to RBD vaccines as well as whole spike vaccines. Data from a challenge study conducted in Syrian hamsters after two doses of RBD-TM Beta mRNA vaccine.

**Figure S8.**
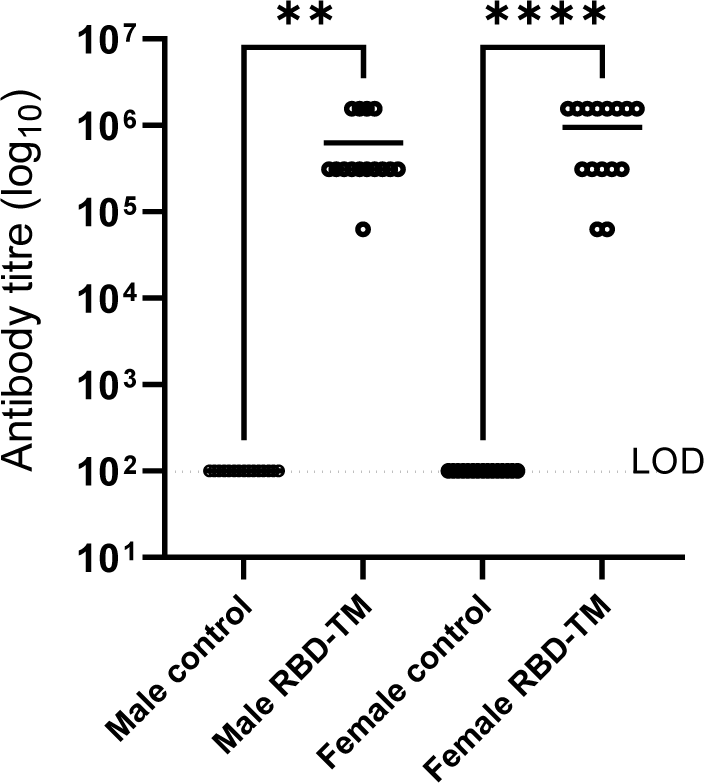
Antibody titres in serum of vaccinated rats. **Background:** Toxicity studies of the Beta RBD-TM mRNA vaccine were carried out to support the proposed Phase 1 clinical trial. The primary purpose of this study was to administer three doses of vaccine to rate using the highest dose proposed for the clinical study. Serum was collected at the end of the study to carry out ELISA analysis of Beta RBD-specific antibodies. **Method:** Male and female Sprague Dawley rats (50% of each sex) were sourced from Animal Resources Centre, Canning Vale WA, Australia. After a 6-day acclimation period, dosing started at 6 weeks of age. 15 rats were used per group. The study was performed at TetraQ, University of Queensland, approved by the University of Queensland Animal Ethics Committee and assigned the following project code: 2021/AE000384. Rats were housed in their study groups up to three rats per cage per sex in an individually ventilated BioZone Global cage system. These solid floor cages are ventilated with filtered air at a minimum of 15 air changes per hour and the rats were provided food ad-libitum and enrichment including red Perspex hutches, aspen chew sticks and kimwipes. **Results:** Rats were monitored daily for any adverse reactions to the treatments. Daily monitoring of the rats included examination for changes in skin and fur, eyes and mucous membranes, respiratory and circulatory function, gait and posture, behaviour, tremors or convulsions and any other abnormal findings. These observations also included daily examination of the injection site for reactions including oedema and erythema. A humane endpoint of 15% body weight loss was established and severe findings in the above categories triggered an immediate inspection by the study veterinarian. No humane endpoints were reached in this study. At the end of the study ELISA data showed that both male and female rats developed high titres of RBD-specific antibodies (Figure 7). Antibody titres measured in rat serum after three 50 μg intramuscular doses of RBD-TM vaccine in the LNP formulation chosen for the Phase 1 clinical study.

**Figure S9:**
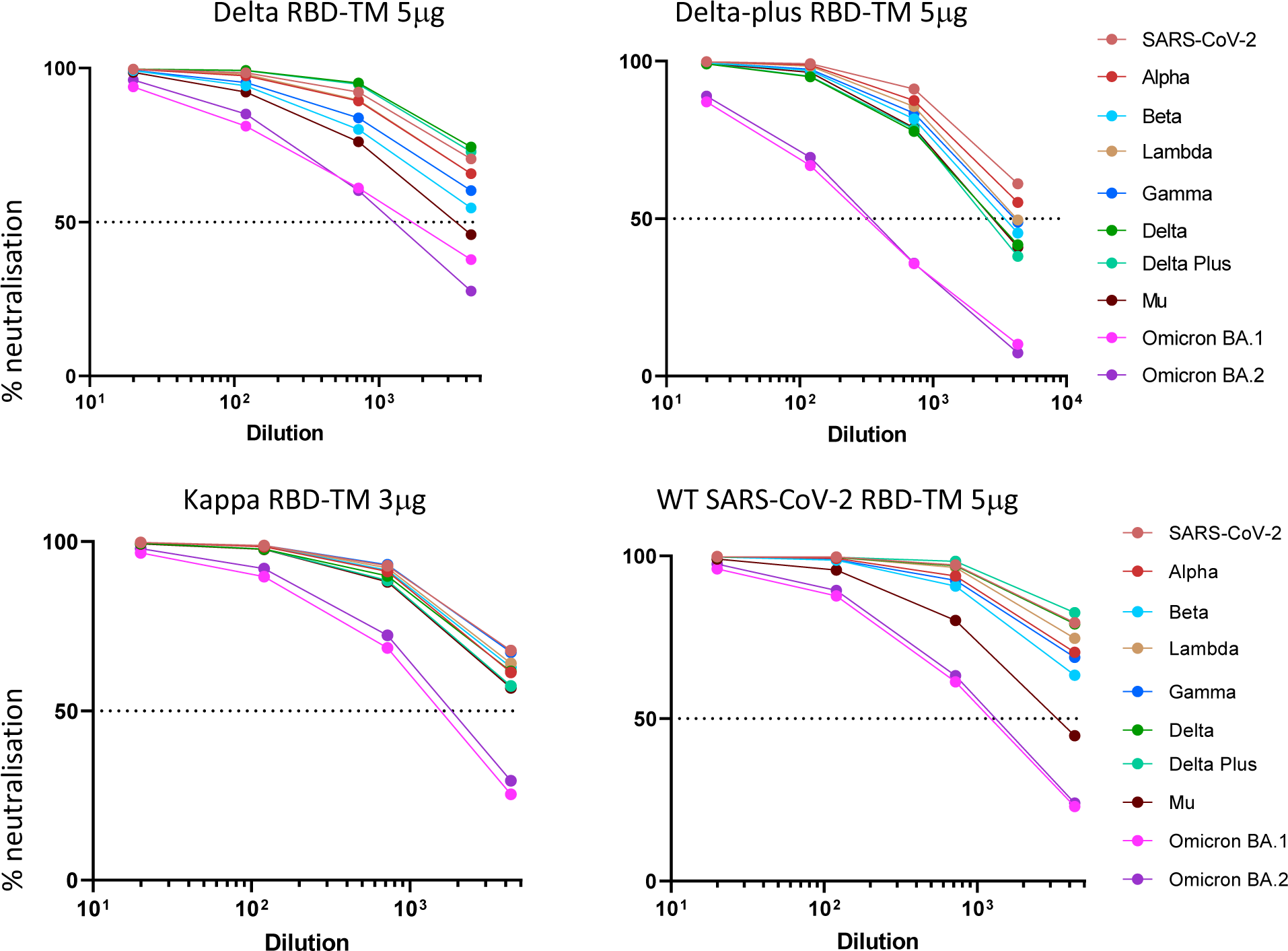
sVNT analysis of mouse serum samples following vaccination of naïve mice with Delta, ‘Delta-plus’ or Kappa RBD-TM vaccines. WT RBD-TM at 5μg is included for comparison. Mice were vaccinated on day 0 and day 21. Graphs show mean % neutralisation of binding of variant RBD-beads to ACE2 plotted against dilution of the mouse serum samples collected on day 56 (n = 5 mice).

**Figure S10:**
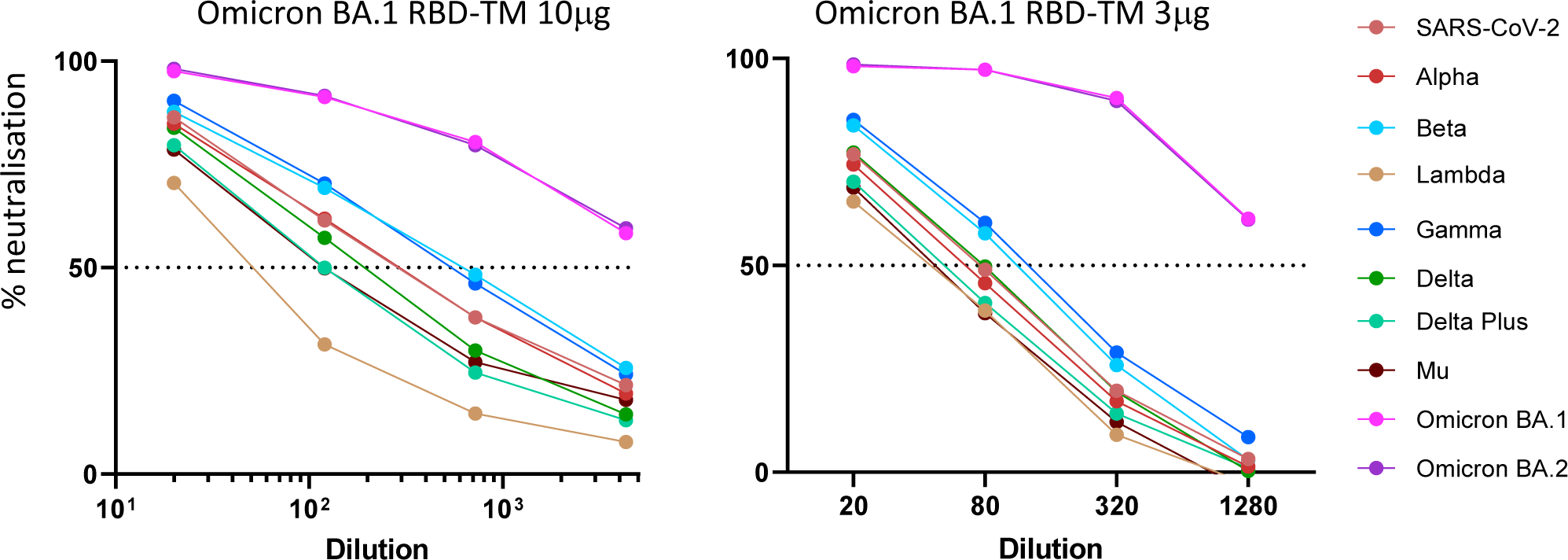
sVNT analysis of mouse serum samples following vaccination of naïve mice with 3μg or 10μg doses of Omicron BA.1 RBD-TM vaccine. Mice were vaccinated on day 0 and day 21. Graphs show mean % neutralisation of binding of variant RBD-beads to ACE2 plotted against dilution of the mouse serum samples collected on day 56 (n = 5 mice).

**Figure S11:**
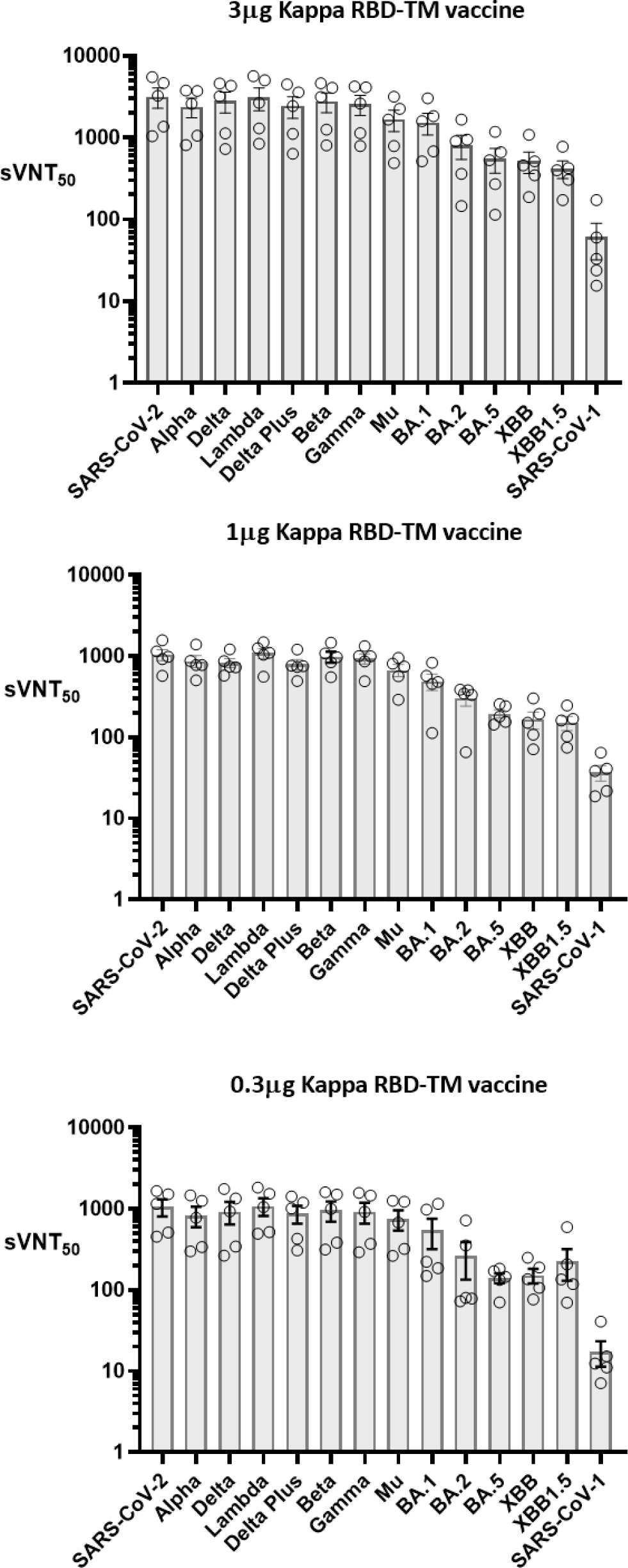
sVNT_50_ values determined using serum samples taken on day 56 from individual mice after vaccination with either 3, 1 or 0.3μg Kappa RBD-TM vaccine. Naïve mice were vaccinated on day 0 and day 21. Open circles represent sVNT_50_ values (mean of two technical replicates) for each mouse. Error bars show the mean and sem sVNT_50_ values (n = 5 mice).

